# Designing multi-epitope based peptide vaccine targeting spike protein SARS-CoV-2 B1.1.529 (Omicron) variant using computational approaches

**DOI:** 10.1101/2021.12.23.473990

**Authors:** Meet Parmar, Ritik Thumar, Jigar Sheth, Dhaval Patel

**Affiliations:** Department of Biological Sciences and Biotechnology, Institute of Advanced Research, Koba Institutional Area, Gandhinagar-382426, Gujarat, India.

**Keywords:** SARS-CoV-2, Vaccine design, Immunoinformatics, Spike protein B1.1.529, Omicron, Multi-subunit epitope vaccine

## Abstract

Since the SARS-CoV-2 outbreak in 2019, millions of people have been infected with the virus, and due to its high human-to-human transmission rate, there is a need for a vaccine to protect people. Although some vaccines are in use, due to the high mutation rate in the SARS-CoV-2 multiple variants, the current vaccines may not be sufficient to immunize people against new variant threats. One of the emerging variants of concern is B1.1.529 (Omicron), which carries∼30 mutations in the Spike protein of SARS-CoV-2 is predicted to evade antibodies recognition even from vaccinated people. We used a structure-based approach along with an epitope prediction server to develop a Multi-Epitope based Subunit Vaccine (MESV) involving SARS-CoV-2 B1.1.529 variant spike glycoprotein. The predicted epitope with better antigenicity and non-toxicity were used for designing and predicting vaccine construct features and structure models. The MESV construct In-silico cloning in pET28a expression vector predicted the construct to be highly translational. The proposed MESV vaccine construct was also subjected to immune simulation prediction and was found to be highly antigenic and elicit a cell-mediated immune response. The proposed MESV in the present study has the potential to be evaluated further for vaccine production against the newly identified B1.1.529 (Omicron) variant of concern.

## Introduction

The world witnessed a fatal outbreak of a novel coronavirus (COVID-19) infection, also known as severe acute respiratory syndrome coronavirus 2 (SARS CoV 2) at the end of the year 2019 (Chan et al., 2020; Wang et al., 2020). Preliminary identified as a causative agent for series of unusual pneumonia in Wuhan City, China and later leading to the sporadic inter-continental pandemic outbreak (Chan et al., 2020; Wu et al., 2020; Zhou et al., 2020). In March 2020, as the infection has crossed most of the international borders, WHO designated the pathogen as SARS-CoV-2 and the disease as coronavirus disease 2019 (COVID-2019) (WHO.int). By January 2022, more than >31.8 million confirmed cases with >5.5 million death reported in 208 countries (WHO.int). The pandemic outbreak affected millions of lives globally due to consequences of mandatory isolations/quarantines and lockdown as well as travel restrictions resulting into economic mayhem (Nicola et al., 2020; *Socio-economic impact of COVID-19 | United Nations Development Programme*, 2020). In humans, COVID-19 infections leads to respiratory tract infections presenting vast range of pathophysiological symptoms ranging from mild, such as fever, coughing, and shortness of breath, and in severe symptoms, such as pneumonia, multiple organ failure (kidney, liver, heart & CNS) and severe acute respiratory failure leading to fatality (Chan et al., 2020; Hui et al., 2020).

The coronaviruses (CoVs) are termed due to crown-like spikes on their surface are classified into four main types α, β, γ and δ infecting wide ranges of hosts ranging from mammals to aves (Cui et al., 2019). The SARS-CoV-2 is a β□coronavirus with a nucleocapsid of helical symmetry, belongs to the subfamily Orthocoronavirinae, in the family Coronaviridae, order Nidovirales (Defay & Cohen, 1995; Zhu et al., 2020). The genome of SARS-CoV-2 is a (+) sense single-stranded RNA as the genetic component which encodes a set of viral replicase proteins, structural proteins including spike S, membrane, envelope, nucleocapsid, proteases which cleaves polyproteins and several uncharacterized proteins (Rota et al., 2003; Zhu et al., 2020). The SARS-CoV-2 genomes encode for 14 ORFs, including ORF1a and ORF1ab which encodes for 16 non-structural proteins (NSP1-16) and four essential structural proteins, including spike (S) glycoprotein, envelope (E) protein, membrane (M) protein and nucleocapsid (N) protein which are vital for viral assembly and invasion of SARS-CoV-2 (Behmard et al., 2020) (Lu et al., 2020). The polyproteins 1a and 1ab (pp1a and pp1ab) are auto-proteolytically cleaved by SARS-CoV-2 proteases to release 11 (pp1a) and 5 (pp1ab) functional proteins necessary for viral replication, survival and proliferation (Báez-Santos et al., 2015; Hoffmann et al., 2020; Ziebuhr et al., 2000). The homotrimer Spike (S) protein constitute the spikes on the viral envelope are responsible for attachment to host cells, the three transmembrane domains consisting M protein determines the virion shape, facilitate membrane curvature, and binds to nucleocapsid, the E protein plays a role in virion assembly, pathogenesis and release, and the N protein is important for binding with the viral RNA genome (Naqvi et al., 2020).

The mode of entry of SARS-CoV-2 is identical to SARS-CoV, using human angiotensin-converting enzyme 2 (ACE2) and CLEC4M/DC-SIGNR receptor for attachment to the cell membrane (Hwang et al., 2020; Walls et al., 2020; Wan et al., 2020). The ACE2 is also found to express on the mucosa of the epithelial cell and the oral cavity of the tongue, type II alveolar cells (AT2), amongst others, acting as entry routes for virus through endosomal pathways (Xu et al., 2020). The SARS-CoV-2 Spike protein (S) is a homotrimer class I type fusion protein that enables viral entry to human cells via ACE2 receptor (Walls et al., 2020). The Spike protein (S) is a 1273 residue glycoprotein with three conformational states: pre-fusion native state, pre-hairpin intermediate state, and post-fusion hairpin state (spike protein) (Hwang et al., 2020). The spike protein is constituted of two subunits, S1 which is required for receptor recognition and S2 required for membrane fusion. The S1 subunit has C-terminal RBD (receptor-binding domain) which primarily interacts with the human ACE2 receptor (Yuan et al., 2020). The S protein undergoes a conformational change when it fuses with the ACE2 receptor destabilizing the pre-fusion trimer, resulting in the discharge of the S1 subunit. This permits the transition of the S2 subunit to a steady post-fusion state (De Wilde et al., 2017). Then the host cell-mediated S protein priming by a cellular serine protease TMPRSS2 is responsible for the virus entering into the host cell (Hoffmann et al., 2020; Hwang et al., 2020). After the virus has entered the host cell, it follows the typical cycle of replication, infection and proliferation of a (+) sense RNA virus such as MERS-CoV and SARS-CoV (Saha et al., 2020).

So far, since the pandemic breakout, the S glycoprotein is known to have a pivotal role in viral attachment and infection, in the induction of neutralizing-antibody and T-cell responses, as well as protective immunity thus designating it as a top candidate for vaccine production. A vaccine based on the spike protein and its epitope could induce antibodies in response to block virus binding and fusion (Ishack & Lipner, 2021; Saha et al., 2020). RNA viruses are known to develop high mutation rates which are related to virus high proliferation, infection and evasion of the host immune system. These high mutation rates are due to the low fidelity of the RNA-dependent RNA polymerase (RdRp) (Duffy, 2018). It also has become evident that the high mutation frequency in the S protein is correlated with specific variants of concern that have caused significant mortality in certain geographical regions of the globe (Rees-Spear et al., 2021; Toyoshima et al., 2020). Since the COVID-19 pandemic outbreak, viral genome sequencing has been accelerated and shared at an unprecedented rate to identify the lineages and trace the virus evolution (*SARS-CoV-2 Variant Classifications and Definitions*, 2021). More than one million SARS-CoV-2 sequences are available via the Global Initiative on Sharing All Influenza Data (GISAID), enabling real-time surveillance of the pandemic unfolding (Meredith et al., 2020). Since late 2020, SARS-CoV-2 evolution has witnessed numerous set of mutations, labelling the new variants as ’*variants of concern’*, that changes the virus characteristics, such as transmissibility and antigenicity (Harvey et al., 2021). A large number of SARS-CoV-2 variants has been recently revealed through genome sequencing projects and the characteristics mutations of these variants are unfortunately the targets of most of the currently licensed and used COVID-19 vaccines (Jia & Gong, 2021). Recent reports have summarized most of the SARS-CoV-2 variants identified and their mutational landscape (Harvey et al., 2021; Jia & Gong, 2021; Lauring & Hodcroft, 2021; *SARS-CoV-2 Variant Classifications and Definitions*, 2021; *Tracking SARS-CoV-2 variants*, 2021; Wahid et al., 2021).

A recent emerging cluster of cases reported in South Africa, identified new variants of SARS-CoV-2 by a team led by Tulio de Oliveira, at the University of KwaZulu-Natal. The variant carries a large number of mutations also found in other variants, including Delta, and reports suggest that it is spreading at warp speed across South Africa and anecdotal reports across other countries too. Due to the high rate of infectivity, WHO designated the strain as B.1.1.529, a variant of concern naming it Omicron on 26 November 2021. The Omicron joins Delta, Alpha, Beta and Gamma on the current WHO list of variants of concern (*Tracking SARS-CoV-2 variants*, 2021). The B.1.1.529 (Omicron) variant harbours a total of 30 amino acid substitutions, three small deletions, and one small insertion in the SARS-CoV-2 spike protein - the host immune system’s prime target that antibodies recognize, potentially dampening their potency.

Out of these 30 mutations, some of them were also present in the other strains such as Delta and Alpha were found to be problematic and linked to high infectivity and enhanced ability to evade antibodies (*Science Brief: Omicron (B.1.1.529) Variant | CDC*, 2021; *Tracking SARS-CoV-2 variants*, 2021). The mutational profile of 30 mutations can be found in table 1. The key amino-acid substitutions in the Spike protein are A67V, del69-70, T95I, del142-144, Y145D, del211, L212I, ins214EPE, **G339D, S371L, S373P, S375F, K417N, N440K, G446S, S477N, T478K, E484A, Q493R, G496S, Q498R, N501Y, Y505H**, T547K, D614G, H655Y, N679K, P681H, N764K, D796Y, N856K, Q954H, N969K, L981F (Substitutions in RBD region (No. 15 mutations) are highlighted in bold type) (Figure 1). The RBD domain of the spike protein is the prime target of neutralizing antibodies generated following infection by SARS-CoV-2 (Liu et al., 2020; Piccoli et al., 2020) and the same is the component of currently used mRNA and adenovirus-based vaccines licensed for use and others awaiting regulatory approval (Dai & Gao, 2020).

**Figure 1:**
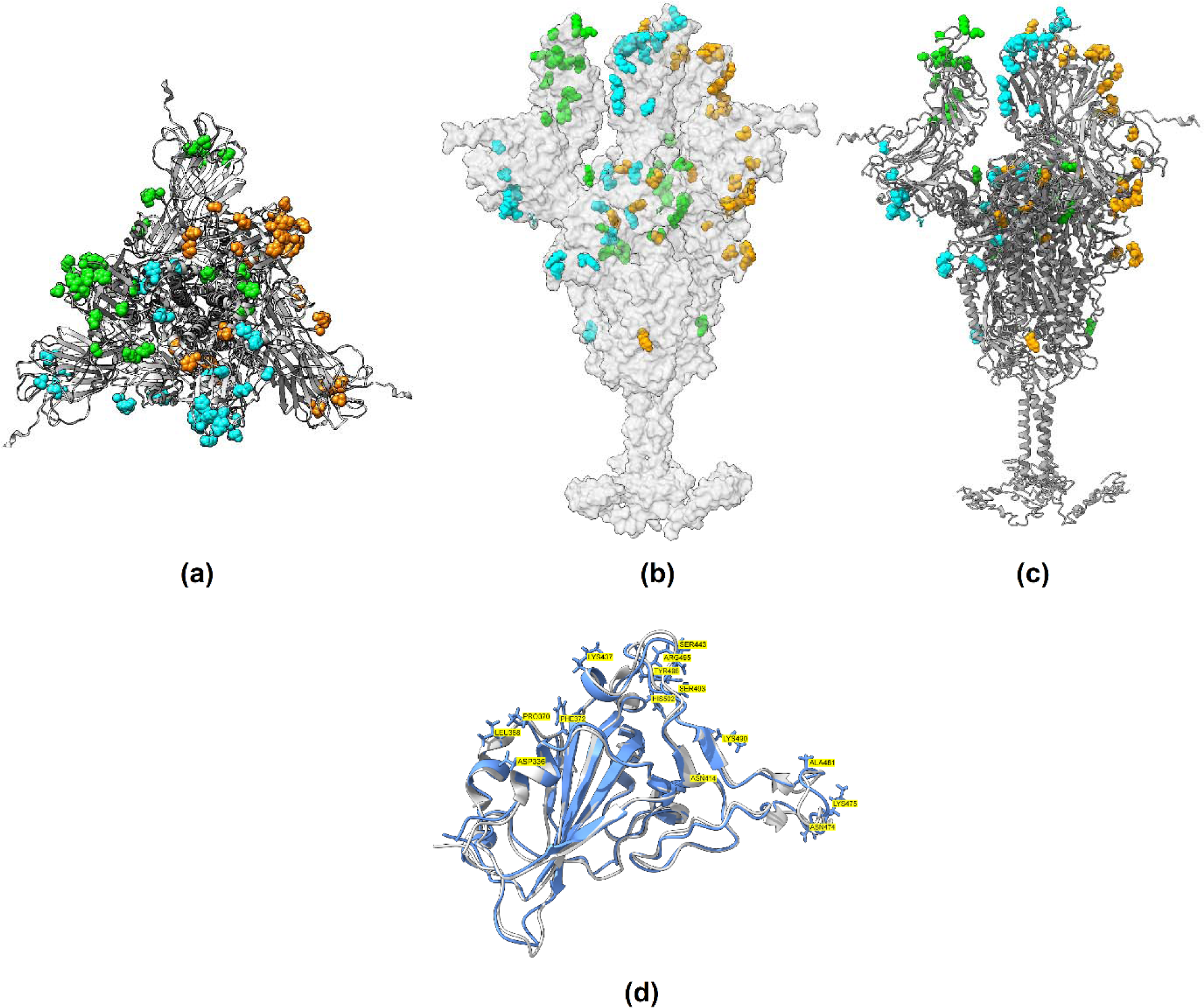
SARS-CoV-2 Spike glycoprotein (B.1.1.529 Omicron) trimer structure model in different representation: top view (a), surface representation (b), cartoon representation (c). The protein chain is colored in grey and mutations in the B.1.1.529 Omicron variant are colored in cyan, orange and green respectively for three chains in trimer model. (d) Structural superimposition of RBD domain from Spike proteins of Wuhan reference (grey) and B.1.1.529 Omicron variant (blue). The mutations in B.1.1.529 Omicron spike protein is labeled with residue and number in yellow box.

**Table 1:**
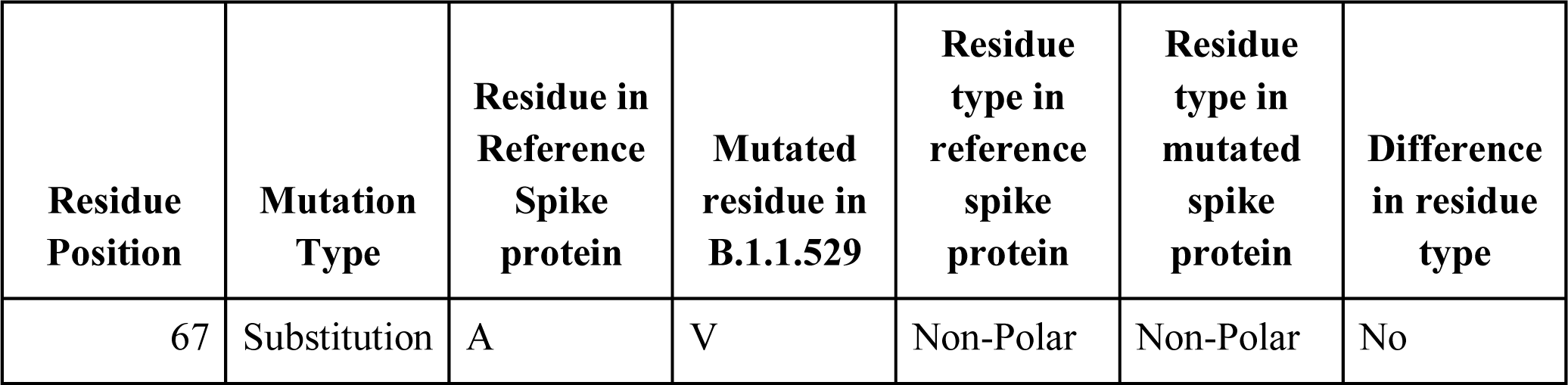
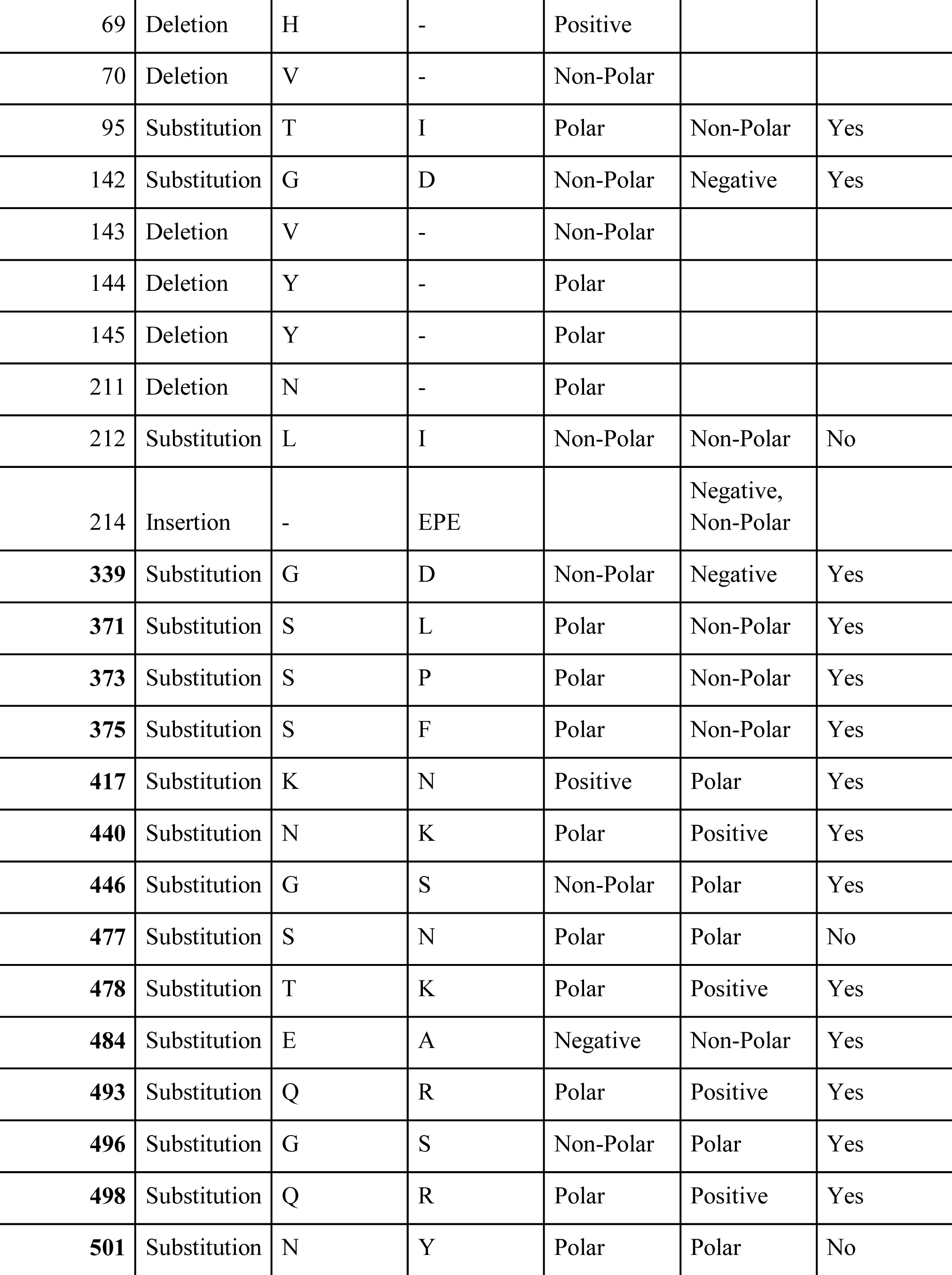
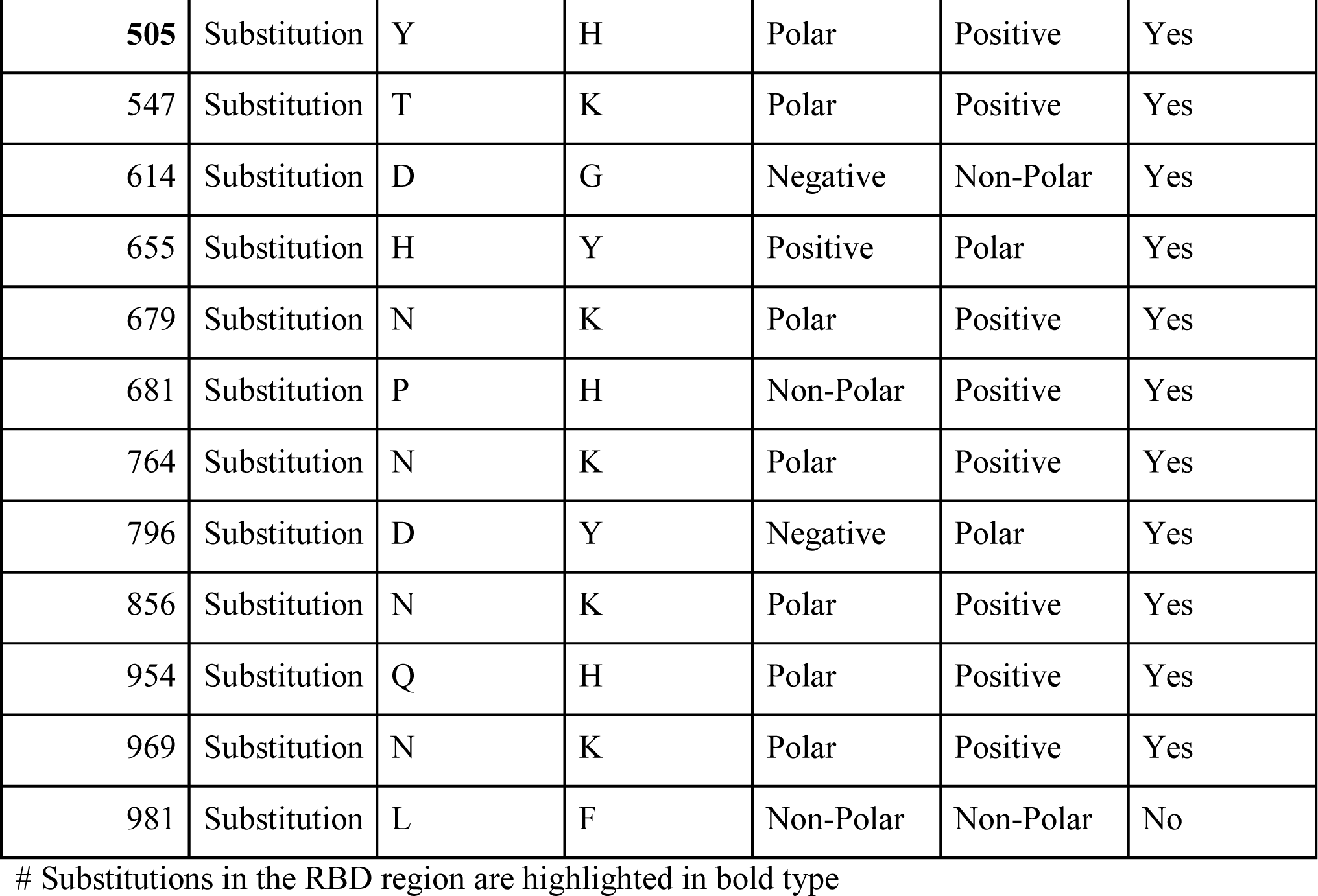
Details of B.1.1.529 variant spike protein mutations, its position, type and difference in comparison to wild-type spike protein.

| Residue Position | Mutation Type | Residue in Reference Spike protein | Mutated residue in B.1.1.529 | Residue type in reference spike protein | Residue type in mutated spike protein | Difference in residue type |
| --- | --- | --- | --- | --- | --- | --- |
| 67 | Substitution | A | V | Non-Polar | Non-Polar | No |
| 69 | Deletion | H | - | Positive |  |  |
| 70 | Deletion | V | - | Non-Polar |  |  |
| 95 | Substitution | T | I | Polar | Non-Polar | Yes |
| 142 | Substitution | G | D | Non-Polar | Negative | Yes |
| 143 | Deletion | V | - | Non-Polar |  |  |
| 144 | Deletion | Y | - | Polar |  |  |
| 145 | Deletion | Y | - | Polar |  |  |
| 211 | Deletion | N | - | Polar |  |  |
| 212 | Substitution | L | I | Non-Polar | Non-Polar | No |
| 214 | Insertion | - | EPE |  | Negative,<br>Non-Polar |  |
| **339** | Substitution | G | D | Non-Polar | Negative | Yes |
| **371** | Substitution | S | L | Polar | Non-Polar | Yes |
| **373** | Substitution | S | P | Polar | Non-Polar | Yes |
| **375** | Substitution | S | F | Polar | Non-Polar | Yes |
| **417** | Substitution | K | N | Positive | Polar | Yes |
| **440** | Substitution | N | K | Polar | Positive | Yes |
| **446** | Substitution | G | S | Non-Polar | Polar | Yes |
| **477** | Substitution | S | N | Polar | Polar | No |
| **478** | Substitution | T | K | Polar | Positive | Yes |
| **484** | Substitution | E | A | Negative | Non-Polar | Yes |
| **493** | Substitution | Q | R | Polar | Positive | Yes |
| **496** | Substitution | G | S | Non-Polar | Polar | Yes |
| **498** | Substitution | Q | R | Polar | Positive | Yes |
| **501** | Substitution | N | Y | Polar | Polar | No |
| **505** | Substitution | Y | H | Polar | Positive | Yes |
| 547 | Substitution | T | K | Polar | Positive | Yes |
| 614 | Substitution | D | G | Negative | Non-Polar | Yes |
| 655 | Substitution | H | Y | Positive | Polar | Yes |
| 679 | Substitution | N | K | Polar | Positive | Yes |
| 681 | Substitution | P | H | Non-Polar | Positive | Yes |
| 764 | Substitution | N | K | Polar | Positive | Yes |
| 796 | Substitution | D | Y | Negative | Polar | Yes |
| 856 | Substitution | N | K | Polar | Positive | Yes |
| 954 | Substitution | Q | H | Polar | Positive | Yes |
| 969 | Substitution | N | K | Polar | Positive | Yes |
| 981 | Substitution | L | F | Non-Polar | Non-Polar | No |

**Table 2:** B-cell linear epitopes of SARS-Cov-2 variant (B.1.1.529) spike protein and their immunogenic properties.

| NO . | Start | End | Peptide | Length | Antigenicity | Toxicity | Allergenicity |
| --- | --- | --- | --- | --- | --- | --- | --- |
| 1 | 1063 | 1078 | HVTYVPAQEKNFTTA<br>P | 16 | 0.8375 | Non-toxic | Non-allergen |
| 2 | 672 | 686 | SYQTQTKSHRRARSV<br>A | 16 | 0.7544 | Non-toxic | Non-allergen |
| 3 | 889 | 903 | AGAALQIPFAMQMAY<br>R | 16 | 0.7318 | Non-toxic | Non-allergen |

Public vaccination is the best approach and tool for controlling and eliminating the pandemic virus, and for the same development of a vaccine is needed urgently (Lu, 2020). Since the outbreak of COVID-19, numerous vaccine candidates are proposed, deployed or under development stage. A total of 139 and 194 vaccine candidates are in clinical and pre-clinical development respectively (WHO.int). Diverse types of vaccine candidates includes but not limited to live attenuated or inactivated whole virus, RNA and DNA based vaccines, subunit vaccines, viral vectors, self-assembling virus-like particle vaccines and others with each having a peculiar advantages and disadvantages. Subunit vaccines constructed on spike protein have been highly effective for eliciting immunogenicity against previous coronavirus break out such as SARS and MERS (Wang et al., 2012; Zhang et al., 2015). One promising approaches includes designing of a subunit vaccine with numerous and diverse antigenic variables which can produce an inclusive spectrum of native viral antigens (Behmard et al., 2020). A typical vaccine development cycle takes a minimum of ten years from lab research to its approved use after all clinical assessments, making such traditional approaches time-consuming and labor intensive (Heaton, 2020; Papaneri et al., 2015). The advent in genome sequencing and high-performance computational technology have made immunoinformatics predictions as cost-effective and convenient (Bahrami et al., 2019; De Groot et al., 2020; Oli et al., 2020). In-silico immuno-informatics approaches have reduced the number of experiments needed for vaccine development and it has demonstrated to identify protein immunogenic attributes with high efficiency (Khan et al., 2014; Naz & Dabir, 2007; Patronov & Doytchinova, 2012).

In this present, we employed immunoinformatics to predict multiple immunogenic proteins from the SARS-CoV-2 proteome and thereby design a multi-epitope vaccine. In the present study, we used computational based immunoinformatics screening approach for the identification and construction of multi-subunit vaccine candidate. We used SARS-CoV-2 spike protein (B.1.1.529 variant) to identify antigenic elements generating both B-cell and T-cell immunity. We predicted B-cell and T-cell epitopes, evaluated its further toxicity, allergenicity and antigenicity properties. We also performed a population coverage analysis and estimated the broad coverage globally. Further B-cell and T-cell epitopes were linked with appropriate adjuvants to generate multi-subunit epitope vaccine (MESV). To estimate the possible immune response generated by identified MESV, we also performed immune simulations. To assess the suitability of recombinant MESV production, we also performed in-silico cloning.

## Materials & Methods

### 1. Data retrieval, structural and physicochemical analysis of SARS-CoV-2 spike protein

The protein sequence of spike protein (B.1.1.529 variant) was derived from the GISAID Database (Accession ID EPI_ISL_6980876). For sequence level comparison with wild type strain, the reference strain (NC 045512.2) was compared with B.1.1.529 variant S protein for sequence similarity using BLAST (Altschul et al., 1990). For tertiary structure prediction of B.1.1.529 S protein, amino-acid sequence was submitted to the I-TASSER (Yang et al., 2014). For structure refinement, the GalaxyRefine (Heo et al., 2013) and ModRefiner (Xu & Zhang, 2011) program was used and to assess the Ramachandran statistics, SAVES server (https://saves.mbi.ucla.edu/) was utilized. The GRAVY (Grand average of hydropathicity), half-life, sub-atomic weight, instability index, aliphatic record, and amino acid atomic composition of the protein sequence were computed through ProtParam (http://web.expasy.org/protparam/).

### 2. Prediction of B-cell linear and discontinuous epitopes

For identification of the linear B cell epitope areas in the antigen sequence, the ABCpred service was utilized. ABCpred operates on the basis of Artificial Neural Networks, which are collections of mathematical models that imitate some of the recognized aspects of the biological nervous system and draw on analogies of adaptive biological learning. The putative epitopes are then tested for antigenicity using Vaxijen 2.0 (Doytchinova & Flower, 2007). We additionally also predicted the discontinuous epitopes that had actually much more substantial attributes compared to the linear epitopes. The identification of discontinuous B-cell epitopes is a significant difficulty in vaccine design. The DiscoTope web server (Kringelum et al., 2012) was used to predict the surface area accessibility as well as amino acids that create discontinuous B-cell epitopes as identified through X-ray crystallography of antigen/antibody protein structures.

### 3. Prediction of CTL and HTL epitopes

The IEDB MHC I binding prediction methods (http://tools.iedb.org/mhci) was used to predict spike glycoprotein CD8+ T cell epitopes from sequence. Using complicated artificial neural network applications, this method unifies epitope prediction restricted to a large number of MHC class I alleles and proteasomal C-terminal cleavage. To predict the CD4+ T cell epitopes (peptides), the MHC II (http://tools.iedb.org/mhcii/) was used. Briefly, the epitopes showing the highest binding diversity with the various HLA serotypes were chosen. Further these epitopes were submitted to Vaxijen 2.0 server for assessing their antigenicity at a 0.7 value threshold. All the top-scoring epitopes identified from each tool mentioned above were then submitted to the IEDB T cell Class I Immunogenicity predictor (http://tools.iedb.org/immunogenicity/). After the determination of the HLA-confined CD8+ and CD4+ T cell epitopes, the epitope toxicity and allergenicity were predicted using ToxinPred server (http://crdd.osdd.net/raghava/toxinpred/) (Gupta et al., 2013) and AlgPred (Saha & Raghava, 2006) server respectively. Finally, the non-allergenic epitopes predicted from the above servers, were chosen as T-cell epitopes.

### 5. Population Coverage analysis

The IEDB Population Coverage tool (http://tools.iedb.org/population/) was used to test chosen epitopes from the HLA class I and class II families, as well as their corresponding binding leukocyte antigens. With a known HLA background, this tool computed the distribution or fraction of persons projected to respond to the selected epitopes. Likewise, the IEDB device processes the normal number of epitope hits/HLA allele blends perceived by the all-out populace, just as the most extreme and least number of epitope hits perceived by 90% of the chosen populace. The HLA genotypic frequencies are estimated, and T cell epitopes are searched by region, ethnicity, and country.

### 6. Designing of multi-epitope vaccine construct

The most efficient epitopes likely to generate subunit immunization should have the qualities such as exceptionally antigenic, immunogenic, non-allergenic, and non-toxic. Those epitopes with the following qualities were chosen further to construct Multi-Epitope Subunit Vaccine (MESV). An adjuvant was also appended with the EAAAK linker to the primary cytotoxic T lymphocytes (CTL) epitope. The different epitopes were connected with AAY, GPGPG, and KK linkers. β-defensin has been used as an adjuvant since it is a 45 amino acids long peptide that goes about as an immunomodulator and as an antimicrobial agent. The 6X His-tag was inserted to the vaccine constructs at the C terminal end for easy purification using affinity chromatography. To check the sequence homology of the MESV construct with human proteome, the MESV sequence was submitted for BLAST sequence homology tool. Physicochemical properties such as like (half-life, hypothetical isoelectric point [pI], hydropathy, and aliphatic index of the MESV construct was assessed using the Protparam tool. The MESV construct’s allergenicity was assessed using the AllerTOP v.2.0 (Dimitrov et al., 2014). For evaluation of the secondary structure component of the MESV construct, PSIPRED (Moffat & Jones, 2021) was used for identification of secondary structural elements such alpha helices, extended chains, the number of beta turns, and random coils.

The homology model of MESV was constructed using I-TASSER (https://zhanggroup.org/I-TASSER/) webserver. The GalaxyRefine (Heo et al., 2013) and ModRefiner (Xu & Zhang, 2011) server were used to energy minimize the predicted MESV 3D structure. The Ramachandran plot statistical evaluation was performed using SAVES webserver, followed by structural validation analysis utilizing the PROSA web server. To estimate the conformational B-cell epitopes of the proposed MESV, the ElliPro tool (http://tools.iedb.org/ellipro/) was used (Ponomarenko et al., 2008). It estimates the residual protrusion index (PI), protein structure, and neighbor residue clustering to predict epitopes.

### 7. Structural modelling, evaluation, and validation

The identified peptides were subjected to structural modeling using PEPFOLD3 (Lamiable et al., 2016) web server. PEP-FOLD is a de-novo method for predicting peptide structures based on amino acid sequences.

### 8. Molecular Docking of the MESV and human immune receptors

Studying the binding affinity of the epitopes, which characterizes their molecular interaction with the human HLA class I and I molecules, is one of the finest approaches to gain insight into their immune response. The CASTp server (http://sts.bioe.uic.edu/castp/) was used to estimate the binding pockets on HLA-class I and II molecules as well as the human immunological receptor (TRL3, PDB: 3ULV & TLR5, PDB: 3J0A). The CASTp server provides extensive and detailed quantitative assessment of a protein’s topographic characteristics. The molecular docking of MESV against TLR3 structure and TLR5 structure (PDB: 3J0A) was carried out using GalaxyPepDock (Lee et al., 2015). The output of the same was re-submitted to ClusPro for further refinement.

### 9. MM/GBSA based evaluation of docked pose

The best scoring dock complex were retrieved from the above docking process and the top pose was assessed through Molecular Mechanics/Generalized Born Surface Area (MM/GBSA) calculations computed by the HawkDock Server (Chen et al., 2016). HawkDock invetigates the docked pose on a per residue basis across electrostatic potentials, Van der Waal potentials, polar solvation free-energy, and solvation free energy using empirical models. The default parameters were used in which the docked pose complex is minimized for 5000 steps, which includes steepest descents (2000 cycles) and conjugate gradient minimizations (3000 cycles) through an implicit solvent model, the ff02 force field.

### 10. In-silico cloning and codon optimization

In order maximize vaccine expression in the host *E. coli* K12 system, codon optimization was carried out using the Java Codon Adaptation Tool (JCat). The restriction enzymes, prokaryotic ribosome binding site, and rho-independent transcription termination were kept in the default state. The acquired codon adaptation index (CAI) value and GC content of the adapted sequence against a certain threshold was assessed. The improved nucleotide was then cloned into the pET28a (+) vector prototype using the SnapGene 4.2 tool.

### 11. MD simulation and Immune Simulation of the Multi Epitope Vaccine

The WAXSIS web-server was utilized for the small-and wide-angle X-ray scattering (SWAXS) to study the properties of the peptide vaccine construct (Knight & Hub, 2015). An accurate prediction of the generated curves was required. The predictions are complicated due to the scattering contribution from the hydration layer and the impact of temperature fluctuations. The MD simulation provides a complete model of the hydration layer and solvent. The protein compactness and the radius of gyration were evaluated by the Guinier analysis. The Vaccine-TLR5 complex was simulated and investigated; the deformability, B factor, eigenvalues, and elastic network of the docked complex were also analyzed.

For evaluation of immunogenicity and immune response characteristics, the MESV was subjected to immune simulation using C-ImmSim online server (http://150.146.2.1/C-IMMSIM/index.php). C-ImmSim predicts related immunological reactions using machine learning approach for three three anatomical compartments: (i) bone, where hematopoietic stem cells are activated and myeloid cells are created, (ii) the lymphatic organ, and (iii) the thymus, where naive T lymphocytes are selected to avoid autoimmune conditions. The simulations were carried out for the administration of two dose injections of MESV at six-week intervals on days 0 and 42. Each time step was positioned at 100 based on the default values, indicating that each time step is 8 hours long and time step 1 is the injection provided at time zero. As a result, two injections were given at six-week intervals. The T cell memory will be evaluated on a constant basis in this situation. The plot analysis was used to generate a graphical interpretation of the Simpson index.

## Result

### 1. Data retrieval, structural and physicochemical analysis of SARS-CoV-2 spike protein

The protein coding sequence of spike protein (B.1.1.529 variant) was retrieved from GISAID database in FASTA format. The sequence homology of spike protein B.1.1.529 variant with reference strain (NC 045512.2) was found to be 97% (Supplementary figure S1). The tertiary structure of full-length spike protein (B.1.1.529 variant) was generated with I-TASSER with a C-score of -1.60 and estimated TM-score of 0.52 ± 0.15. The Ramachandran statistics of 3D model was 73.4% in core, 20.4% in allowed, 3.5% in generously allowed and 2.7% in disallowed region (Figure 2). After rounds of energy refinement, the Ramachandran statistics were improved to 86.7% in core, 9.7% in allowed, 1.6% in generously allowed and 1.9% in disallowed region. The physiochemical attributes of the spike protein (B.1.1.529 variant) are as follow: No. of amino acids (1270), Mol. Wt. 141300 Da, pI 7.14, estimated half-life >10 hours in prokaryotes and 30 hours in eukaryotes, instability index 34.12, aliphatic index 84.95 and GRAVY score of -0.080.

**Figure 2:**
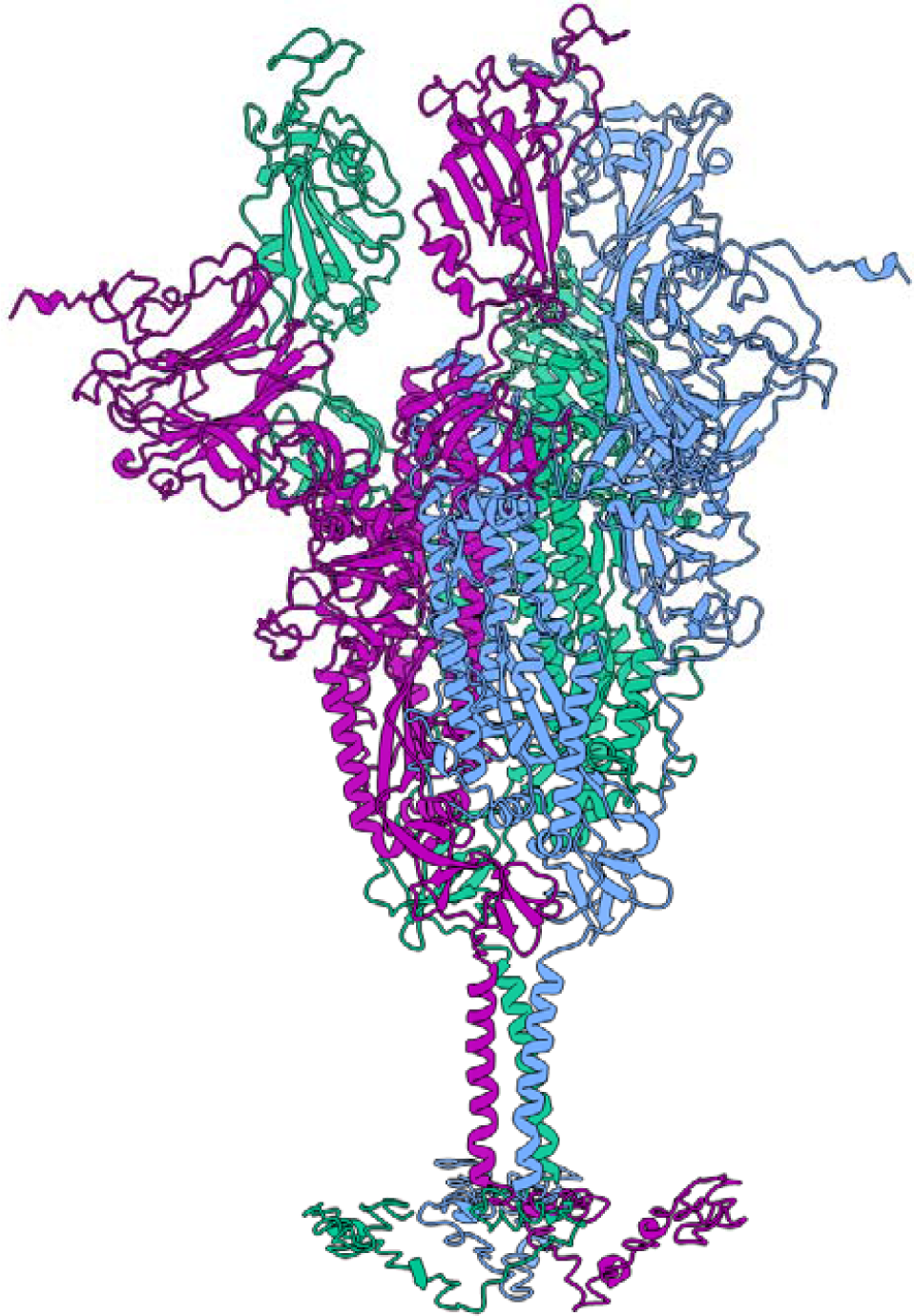
SARS-CoV-2 Spike glycoprotein (B.1.1.529 Omicron) trimer structure model with each color representing three different chains.

### 2. Linear and discontinuous B cell epitopes

B-cell epitopes play a significant role in viral infection resistance. The amino acid screening methods were used to analyze possible B-cell epitopes from the spike protein (B.1.1.529 variant) sequence. To predict possible B-cell epitopes, a consensus-based technique was utilized. Three potential linear B cell epitopes with non-allergenic, non-toxicity and antigenic properties were identified after stringent criteria selection (Table 1). The peptide “HVTYVPAQEKNFTTAP” exhibited the greatest antigenic index when compared to the other predicted B-cell epitope candidates. The linear B-cell epitopes were mapped on the trimeric spike glycoprotein model as shown in figure 3. The discontinuous B-cell epitopes were predicted by Discotope 2.0 using trimeric S protein (B.1.1.529) model. The positions of discontinuous epitopes were mapped on the surface of model structure of S protein (Figure 4). All the discontinuous B-cell epitopes were mapped on the fully-exposed ‘spike head’ region (Figure 4) and no discontinuous B-cell epitopes were found on less-exposed ‘spike stem’ and the ‘spike root’ region of the spike glycoprotein (Figure 4). Predicted discontinuous epitopes were selected from the complete protein chain component of Omicron’s spike glycoprotein and graded according to their propensity score.

**Figure 3:**
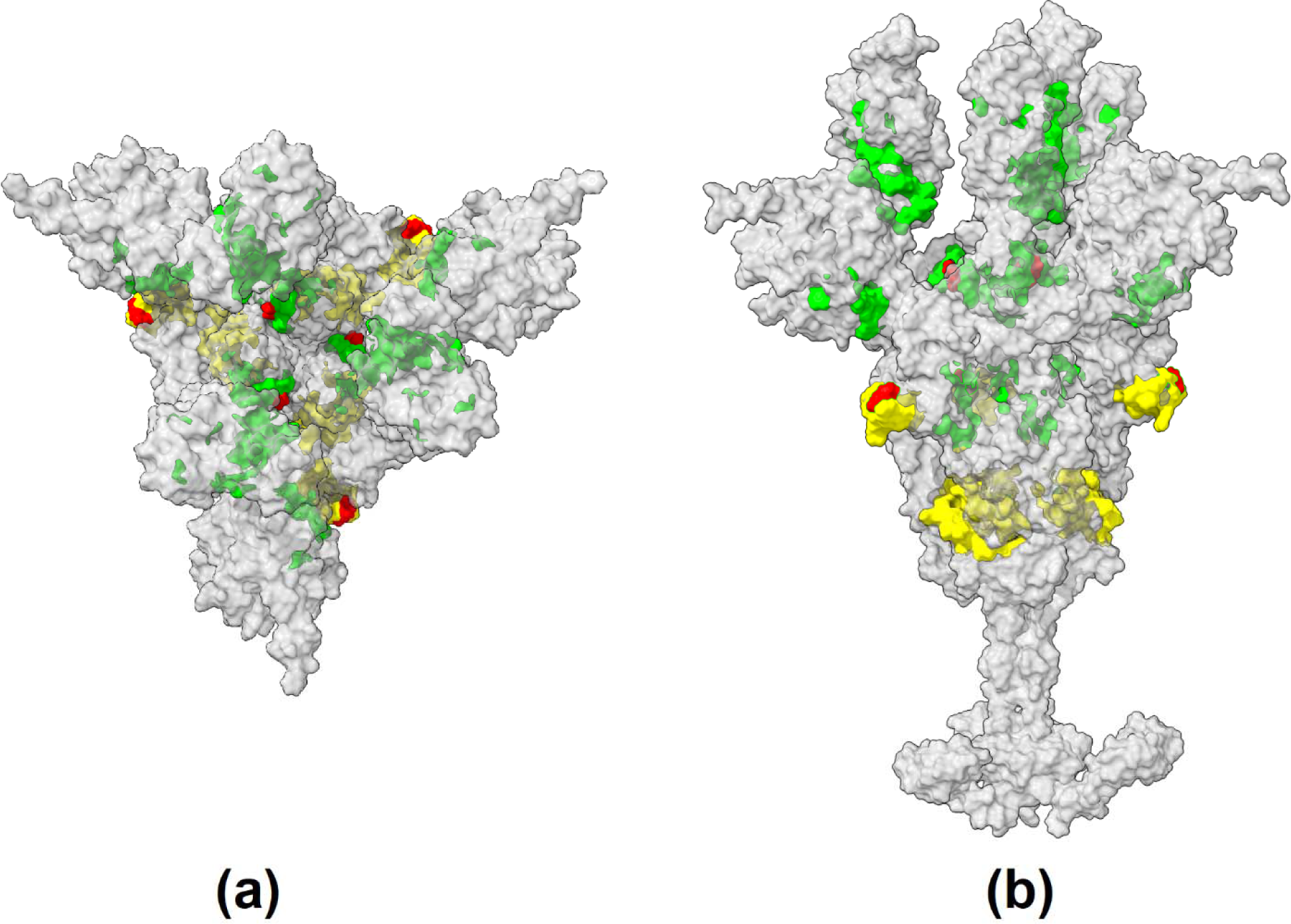
SARS-CoV-2 Spike glycoprotein (B.1.1.529 Omicron) trimer structure model with mapped B-cell (yellow) and T-cell (green) predicted epitopes. The novel mutations of B.1.1.529 Omicron variant in the predicted epitopes are marked in red.

**Figure 4:**
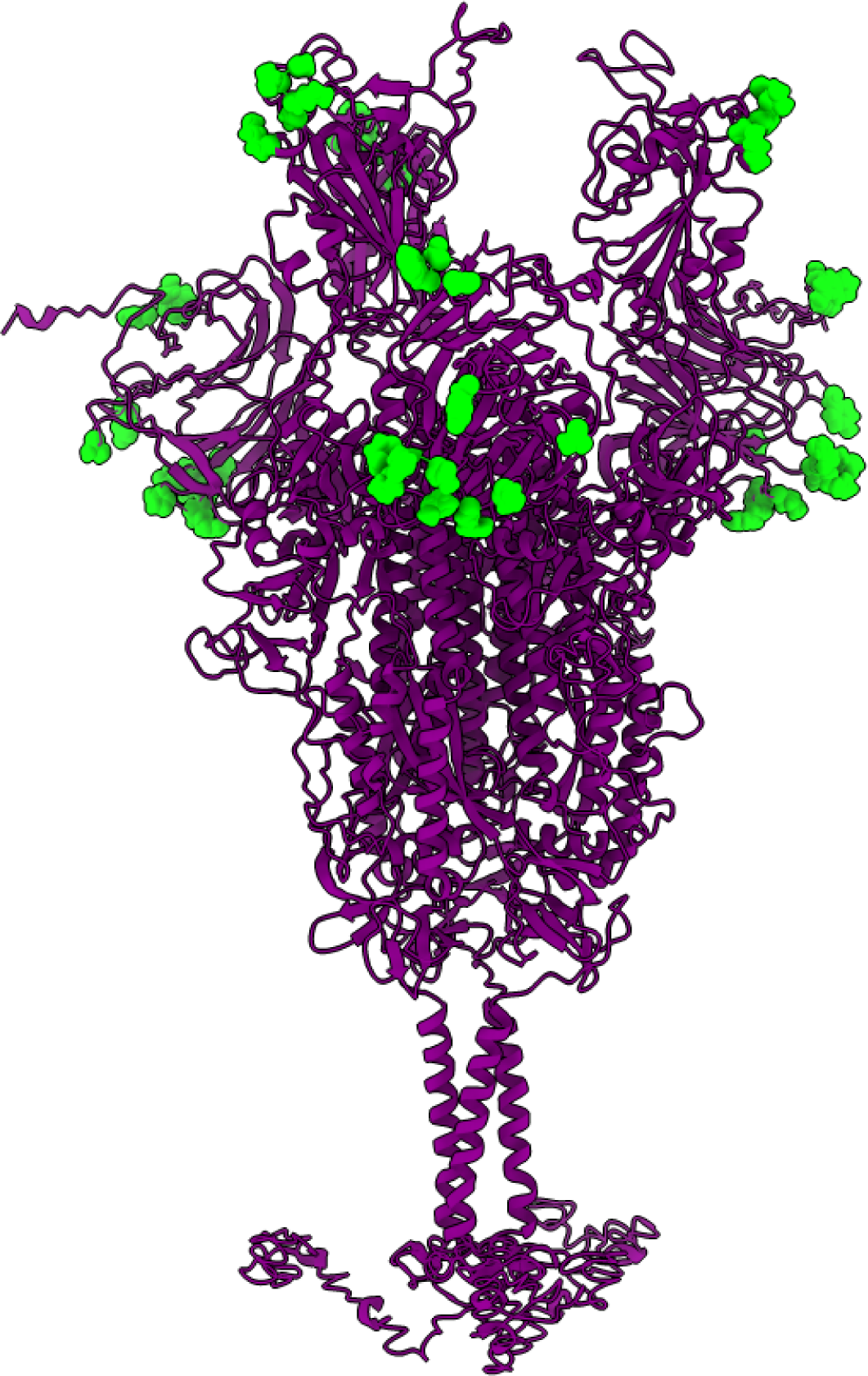
SARS-CoV-2 Spike glycoprotein (B.1.1.529 Omicron) trimer structure model with mapped discontinuous B-cell (green) epitopes.

### 3. T-cell (T_HTL_, and T_CTL_) epitope prediction

The spike (B.1.1.529) protein sequence was tested against several HLA class 1 alleles. The peptides were chosen depending on their rate rankings and the quantity of alleles they possibly bind. Also, the peptides were exposed to antigenicity tests utilizing Vaxijen 2.0. In light of the antigenicity scores, 9 epitopes were chosen for the subsequent analysis. The main peptides are those which have high restricting energy to HLA class I atoms and showing a non-allergenic property. Before considering vaccine design, allergenicity prediction is critical, as vaccination candidates may induce a Type II hypersensitivity reaction. Allergen 1.0 and ToxinPred were used to assess the epitopes non-allergenicity and non-toxicity properties. The spike (B.1.1.529) protein sequence was also searched through a large number of MHC-II alleles for the HLA class II T-cell epitopes. Three epitopes were chosen for their antigenic characteristics. All of these non-allergenic epitopes can elicit an immunological response by activating either or all of the IFN-γ and IL-4 cytokines. An ideal peptide anticipated as epitope should have both high antigenicity as well as capacity to bind with large number of alleles, thus having a high potential to initiate a strong defense response upon immunization. A total of 9 MHC class-I allele binding peptides and 3 MHC class-II allele binding peptides were obtained by using above stringent criteria (Table 4A, 4B). The T-cell epitopes (MHC class I & II) were mapped on the trimeric spike glycoprotein model as shown in figure 3.

The non-allergenic peptides were: “IPFAMQMAY” restricted to HLA-B*35:01,HLA-B*53:01,HLA-B*51:01 HLA allele, “LPFFSNVTW” restricted to HLA-B*35:01,HLA-B*51:01, HLA-B*57: 01,HLA-B*07:02,HLA-B*58:01 with an antigenicity score of 1.0808, “SVYAWNRKR” attaches to 7 alleles HLA-A*31:01, HLA-A*33:01,HLA-A*68:01,HLA-A*03:01,HLA-A*68:01,HLA-A*11:01,HLA-A*30:01, “EILPVSMTK” restricted to HLA-A*68:01,HLA-A*11:01,HL-A*11:01, HLA-A*03:01,HLA-A*33:01,HLA-A*33:01, the peptide “TEILPVSMTK” would be able to bind with 3 alleles HLA-A*11:01, HLA-A*03:01,HLA-A*68:01,”PFAMQMAYR” restricted to HLA-A*33:01, “IPFAMQMAYR” restricted to HLA-A*33:01,HLA-A*35:01, “KFGAISSVL” restricted to HLA-A*24:02,HLA-A*23:01,”PFAMQMAYRF” restricted to HLA-A*23:01,HLA-A*24:02 (Table 3).

**Table 3:** Discontinous B-cell epitope contact numbers and their propensity score in the Spike glycoprotein (B.1.1.529 Omicron).

| Chain ID | Residue ID | Residue Name | Contact Number | Propensity Score | Discotope score |
| --- | --- | --- | --- | --- | --- |
| A | 177 | LYS | 3 | 3.153 | 2.445 |
| A | 497 | THR | 4 | 3.145 | 2.324 |
| A | 442 | VAL | 0 | 2.231 | 1.974 |
| A | 1147 | GLU | 5 | 2.832 | 1.931 |
| A | 178 | GLN | 0 | 1.833 | 1.622 |
| A | 443 | SER | 5 | 2.339 | 1.495 |
| A | 1150 | ASP | 6 | 2.463 | 1.489 |
| A | 1151 | LYS | 6 | 2.397 | 1.432 |
| A | 1155 | ASN | 6 | 2.363 | 1.402 |
| A | 1152 | TYR | 5 | 2.212 | 1.382 |
| A | 1148 | GLU | 6 | 2.291 | 1.338 |
| A | 1154 | LYS | 6 | 2.243 | 1.295 |
| A | 1153 | PHE | 6 | 2.1 | 1.169 |
| A | 1143 | ASP | 6 | 2.019 | 1.097 |
| A | 1146 | LYS | 6 | 1.96 | 1.044 |
| A | 499 | GLY | 0 | 1.174 | 1.039 |
| A | 496 | PRO | 5 | 1.767 | 0.989 |
| A | 1149 | LEU | 7 | 1.991 | 0.957 |
| A | 1156 | HIS | 6 | 1.724 | 0.836 |
| A | 175 | GLU | 1 | 1.061 | 0.824 |
| A | 1145 | PHE | 5 | 1.172 | 0.462 |
| A | 1144 | SER | 6 | 1.193 | 0.366 |
| A | 807 | SER | 6 | 1.172 | 0.347 |
| A | 1141 | GLU | 5 | 0.972 | 0.285 |
| A | 1142 | LEU | 5 | 0.767 | 0.103 |
| A | 1159 | PRO | 3 | 0.426 | 0.032 |
| A | 495 | ARG | 8 | 1.046 | 0.006 |
| A | 1158 | SER | 6 | 0.775 | -0.004 |
| A | 1140 | PRO | 6 | 0.639 | -0.124 |

**Table 4A:**
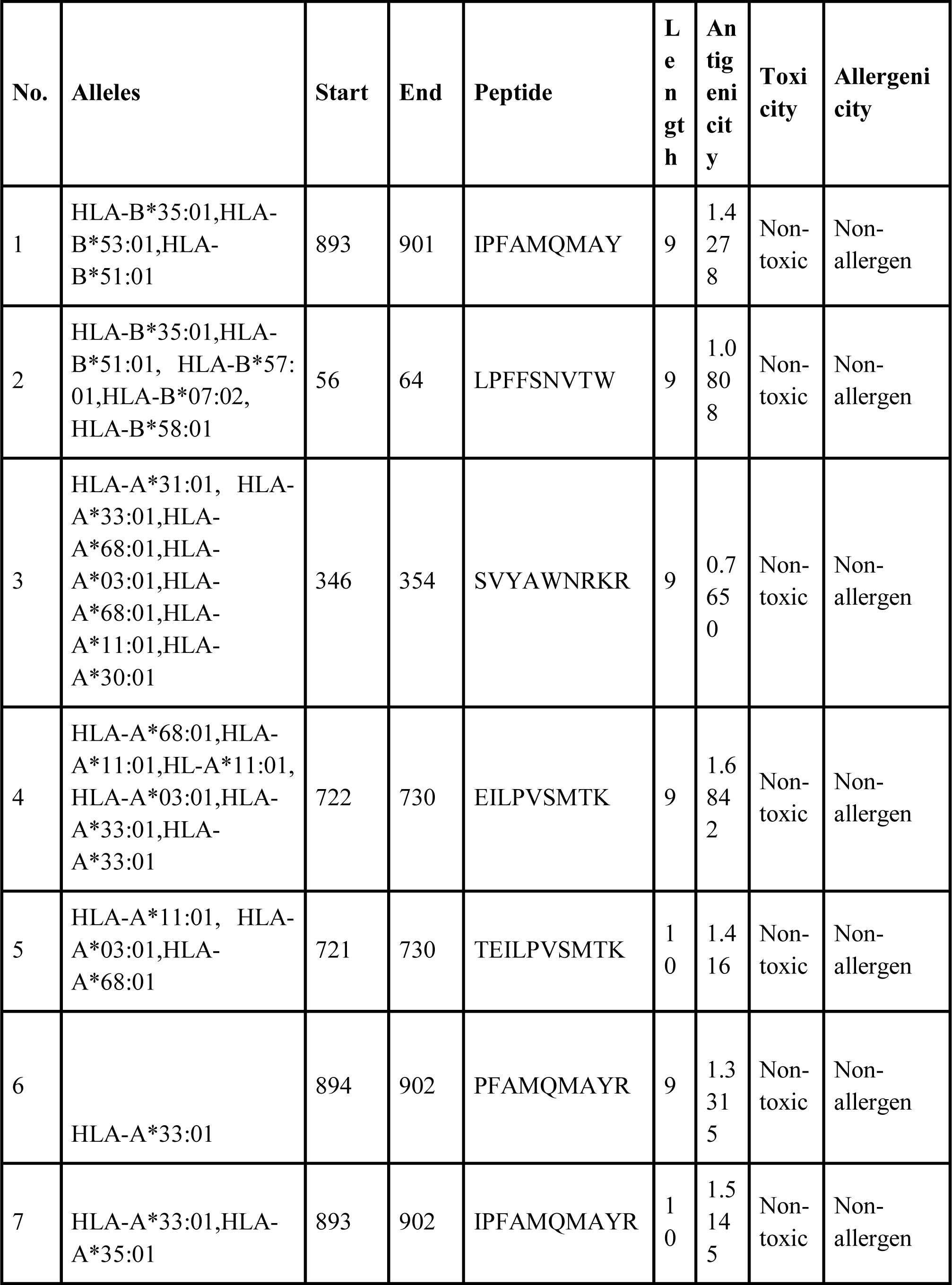
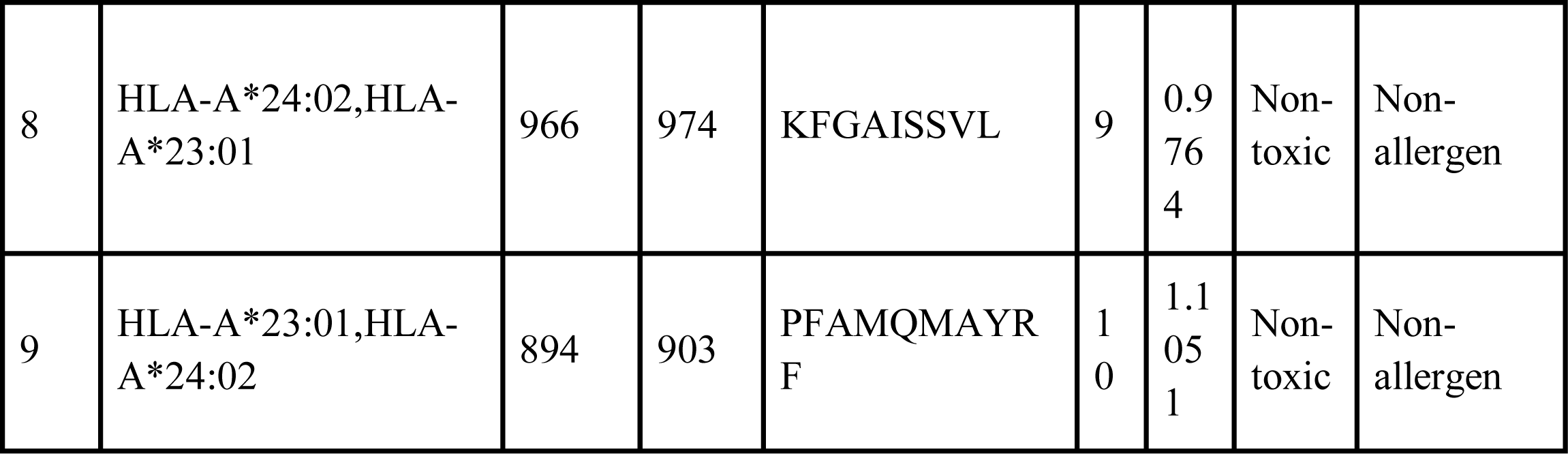
MHC CLASS I Epitopes of SARS-CoV-2 (B.1.1.529) Spike protein.

| No. | Alleles | Start | End | Peptide | Length | Antigenicity | Toxicity | Allergenicity |
| --- | --- | --- | --- | --- | --- | --- | --- | --- |
| 1 | HLA-B*35:01,HLA-B*53:01,HLA-B*51:01 | 893 | 901 | IPFAMQMAY | 9 | 1.4278 | Non-toxic | Non-allergen |
| 2 | HLA-B*35:01,HLA-B*51:01, HLA-B*57:01,HLA-B*07:02,HLA-B*58:01 | 56 | 64 | LPFFSNVTW | 9 | 1.0808 | Non-toxic | Non-allergen |
| 3 | HLA-A*31:01, HLA-A*33:01,HLA-A*68:01,HLA-A*03:01,HLA-A*68:01,HLA-A*11:01,HLA-A*30:01 | 346 | 354 | SVYAWNRKR | 9 | 0.7650 | Non-toxic | Non-allergen |
| 4 | HLA-A*68:01,HLA-A*11:01,HL-A*11:01,HLA-A*03:01,HLA-A*33:01,HLA-A*33:01 | 722 | 730 | EILPVSMTK | 9 | 1.6842 | Non-toxic | Non-allergen |
| 5 | HLA-A*11:01, HLA-A*03:01,HLA-A*68:01 | 721 | 730 | TEILPVSMTK | 10 | 1.416 | Non-toxic | Non-allergen |
| 6 | HLA-A*33:01 | 894 | 902 | PFAMQMAYR | 9 | 1.3315 | Non-toxic | Non-allergen |
| 7 | HLA-A*33:01,HLA-A*35:01 | 893 | 902 | IPFAMQMAYR | 10 | 1.5145 | Non-toxic | Non-allergen |
| 8 | HLA-A*24:02,HLA-A*23:01 | 966 | 974 | KFGAISSVL | 9 | 0.9764 | Non-toxic | Non-allergen |
| 9 | HLA-A*23:01,HLA-A*24:02 | 894 | 903 | PFAMQMAYRF | 10 | 1.1051 | Non-toxic | Non-allergen |

**Table 4B:** MHC CLASS II Epitopes of SARS-Cov-2 (B.1.1.529) Spike protein. [+ induced, -not induced, Allergenicity at a 0.7 threshold].

| No. | Alleles | Start | End | Peptide | Length | Antigenicity | Toxicity | Allergenicity | IFN-γ | IL4 | IL10 |
| --- | --- | --- | --- | --- | --- | --- | --- | --- | --- | --- | --- |
| 1 | HLA-DRB1*01:01,HLA-DRB1*03:01,HLA-DRB1*04:01,HLA-DRB1*04:05,HLA-DRB1*07:01,HLA-DRB1*08:02,HLA-DQA1*03:01/DQB1*03:02,HLA-DQA1*04:01/DQB1*04:02,HLA-DQA1*05:01/DQB1*02:01 | 504 | 518 | PYRVV<br>VLSFEL<br>LHAP | 15 | 0.7072 | Non-toxin | Non-allergen | + | + | - |
| 2 | HLA-DRB1*01:01,HLA<br>-<br>DRB1*03:01,HLA<br>-<br>DRB1*04:01,HLA<br>-<br>DRB1*04:05,HLA<br>-<br>DRB1*07:01,HLA<br>-DRB1*08:02,<br>HLA-DRB4*01:01,HLA<br>-DRB5*01:01 | 50<br>6 | 5<br>2<br>0 | RVVVL<br>SFELLH<br>APAT | 16 | 0.7485 | Non-toxin | Non-<br>allerge<br>n | + | + | - |
| 3 | HLA-DPA1*03:01/DPB<br>1*04:02,HLA-DPA1*02:01/DPB<br>1*01:01,HLA-DPA1*01:03/DPB<br>1*02:01 | 50<br>5 | 5<br>1<br>9 | YRVVV<br>LSFELL<br>HAPA | 15 | 0.7072 | Non-toxin | Non-<br>allerge<br>n | + | + | - |

**Table 4A:**
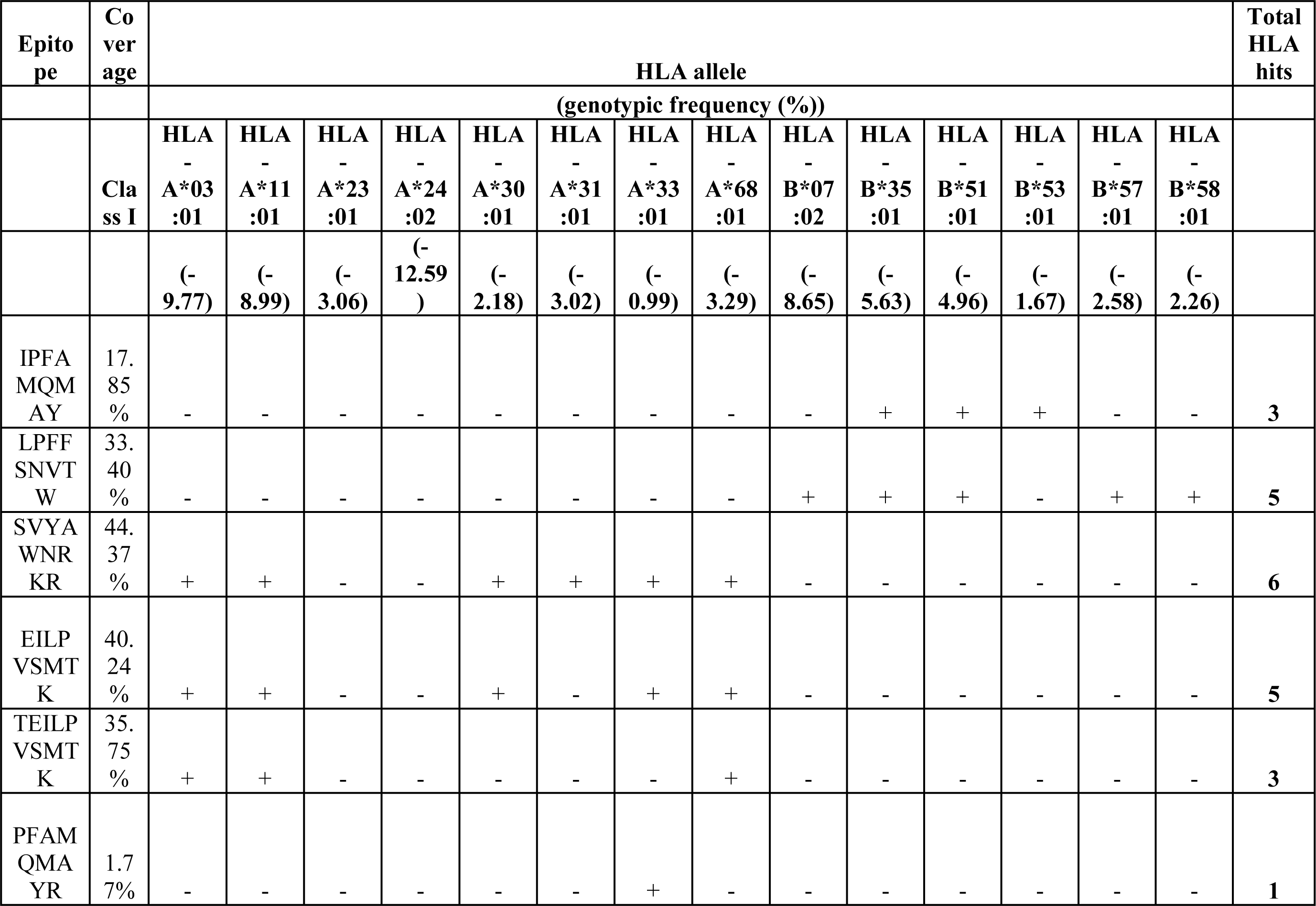
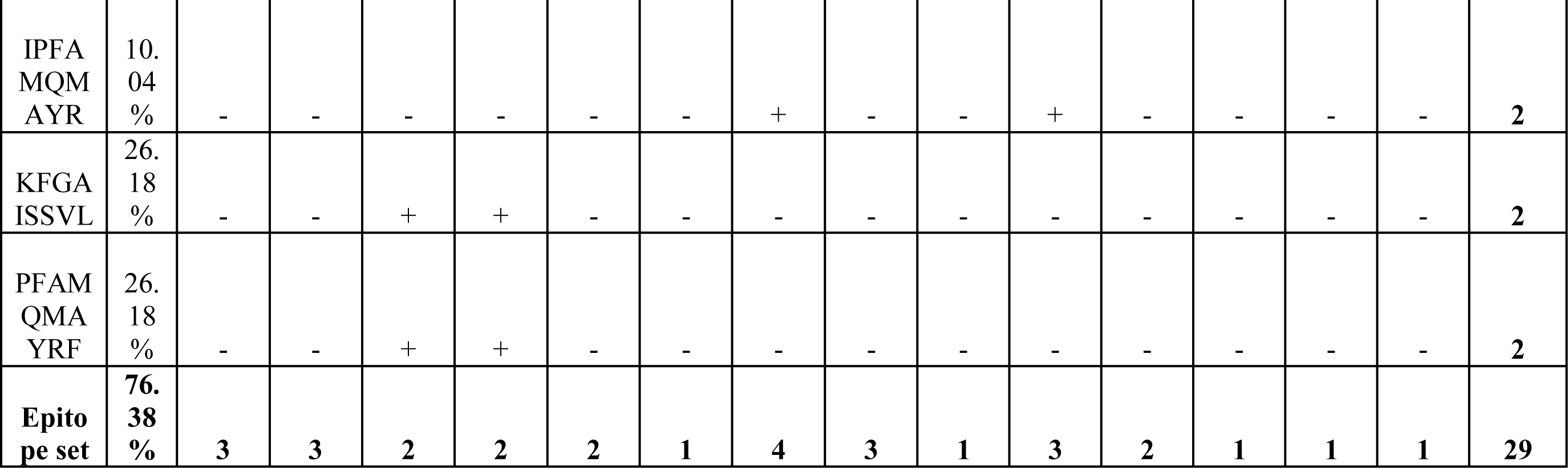
Population coverage analysis of MHC-I class epitopes (+ restricted, − not restricted)

| Epitope | Coverage | HLA allele |  |  |  |  |  |  |  |  |  |  |  |  |  | Total HLA hits |
| --- | --- | --- | --- | --- | --- | --- | --- | --- | --- | --- | --- | --- | --- | --- | --- | --- |
|  |  | (genotypic frequency (%)) |  |  |  |  |  |  |  |  |  |  |  |  |  |  |
|  | Class I | HLA - A*03:01 | HLA - A*11:01 | HLA - A*23:01 | HLA - A*24:02 | HLA - A*30:01 | HLA - A*31:01 | HLA - A*33:01 | HLA - A*68:01 | HLA - B*07:02 | HLA - B*35:01 | HLA - B*51:01 | HLA - B*53:01 | HLA - B*57:01 | HLA - B*58:01 |  |
|  |  | (- 9.77) | (- 8.99) | (- 3.06) | (- 12.59) | (- 2.18) | (- 3.02) | (- 0.99) | (- 3.29) | (- 8.65) | (- 5.63) | (- 4.96) | (- 1.67) | (- 2.58) | (- 2.26) |  |
| IPFA MQM AY | 17.85 % | - | - | - | - | - | - | - | - | - | + | + | + | - | - | 3 |
| LPFF SNVT W | 33.40 % | - | - | - | - | - | - | - | - | + | + | + | - | + | + | 5 |
| SVYA WNR KR | 44.37 % | + | + | - | - | + | + | + | + | - | - | - | - | - | - | 6 |
| EILP VSMT K | 40.24 % | + | + | - | - | + | - | + | + | - | - | - | - | - | - | 5 |
| TEILP VSMT K | 35.75 % | + | + | - | - | - | - | - | + | - | - | - | - | - | - | 3 |
| PFAM QMA YR | 1.77 % | - | - | - | - | - | - | + | - | - | - | - | - | - | - | 1 |
| IPFA<br>MQM<br>AYR | 10.04<br>% | - | - | - | - | - | - | + | - | - | + | - | - | - | - | 2 |
| KFGA<br>ISSVL | 26.18<br>% | - | - | + | + | - | - | - | - | - | - | - | - | - | - | 2 |
| PFAM<br>QMA<br>YRF | 26.18<br>% | - | - | + | + | - | - | - | - | - | - | - | - | - | - | 2 |
| Epito<br>pe set | 76.38<br>% | 3 | 3 | 2 | 2 | 2 | 1 | 4 | 3 | 1 | 3 | 2 | 1 | 1 | 1 | 29 |

**Table 4B:**
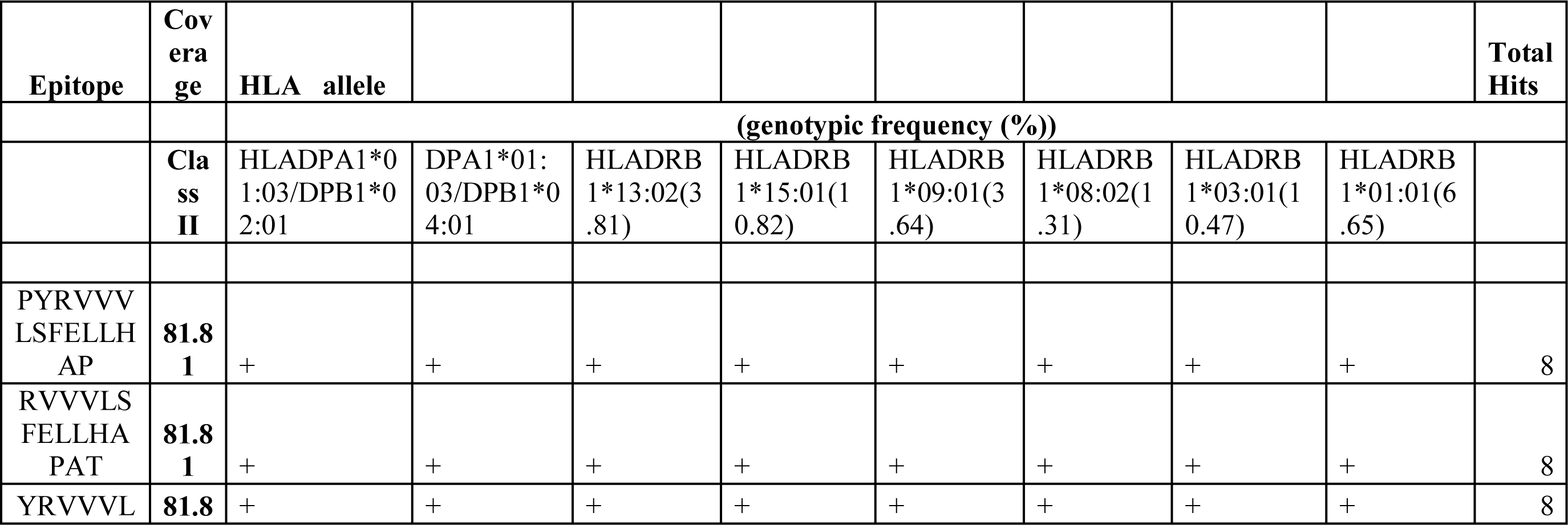
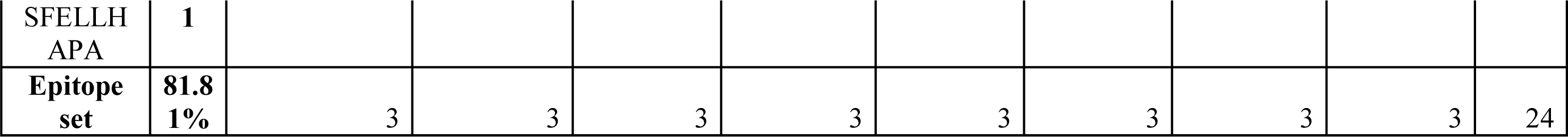
Population coverage analysis of MHC-II class epitopes (+ restricted)

The peptide “PYRVVVLSFELLHAP”, with an antigenicity score of 0.7072, attaches to 8 HLA alleles: HLA-DRB1*01:01, HLA-DRB1*03:01, HLA-DRB1*04:01, HLA-DRB1*04:05, HLA-DRB1*07:01, HLA-DRB1*08:02, HLA-DRB4*01:01, HLA-DRB5*01:01. The epitope “RVVVLSFELLHAPAT”, with an antigenicity score of 0.7485, is also restricted to 8 HLA alleles: HLA-DRB1*01:01, HLA-DRB1*03:01, HLA-DRB1*04:01, HLA-DRB1*04:05, HLA-DRB1*07:01, HLA-DRB1*08:02, HLA-DQA1*03:01/DQB1*03:02, HLA-DQA1*04:01/DQB1*04:02, HLA-DQA1*05:01/DQB1*02:01. The peptide “YRVVVLSFELLHAPA”, binds to allele HLA-DPA1*03:01/DPB1*04:02, HLA-DPA1*02:01/DPB1*01:01, HLA-DPA1*01:03/DPB1*02:01 with the antigenicity score of 0.7072. The HLA-DRB1 is the most common and versatile MHC-II molecule.

### 4. Population Coverage of CTL and HTL epitopes

The distribution of HLA alleles varies among heterogeneous ethnic groups and geographical regions. Therefore, while constructing a viable epitope based vaccine, it is necessary to take population coverage in consideration. When examined with the entire global population, the selected MHC-I class epitopes showed higher individual percentage cover. The 76.38% of the world individuals are capable of responding to the median of one MHC-I epitope. (Table 4A)

It was inferred that, out of the selected continents and subcontinents, North America would possibly show a significant response to the selected HLA-I class restricted epitopes. East Asia, Oceania, West Indies, West Africa, Northeast Asia and Southeast Asia had the highest population coverage of 83.65%, 82.01%, 81.85%, 79.54%, 77.90%, 77.45% respectively, while the Central America population had the lowest population coverage at 6.44%. The MHC-I class epitopes population coverage has been considerably higher as opposed to the MHC-II class epitopes (Table 4B) (Figure 5).

**Figure 5:**
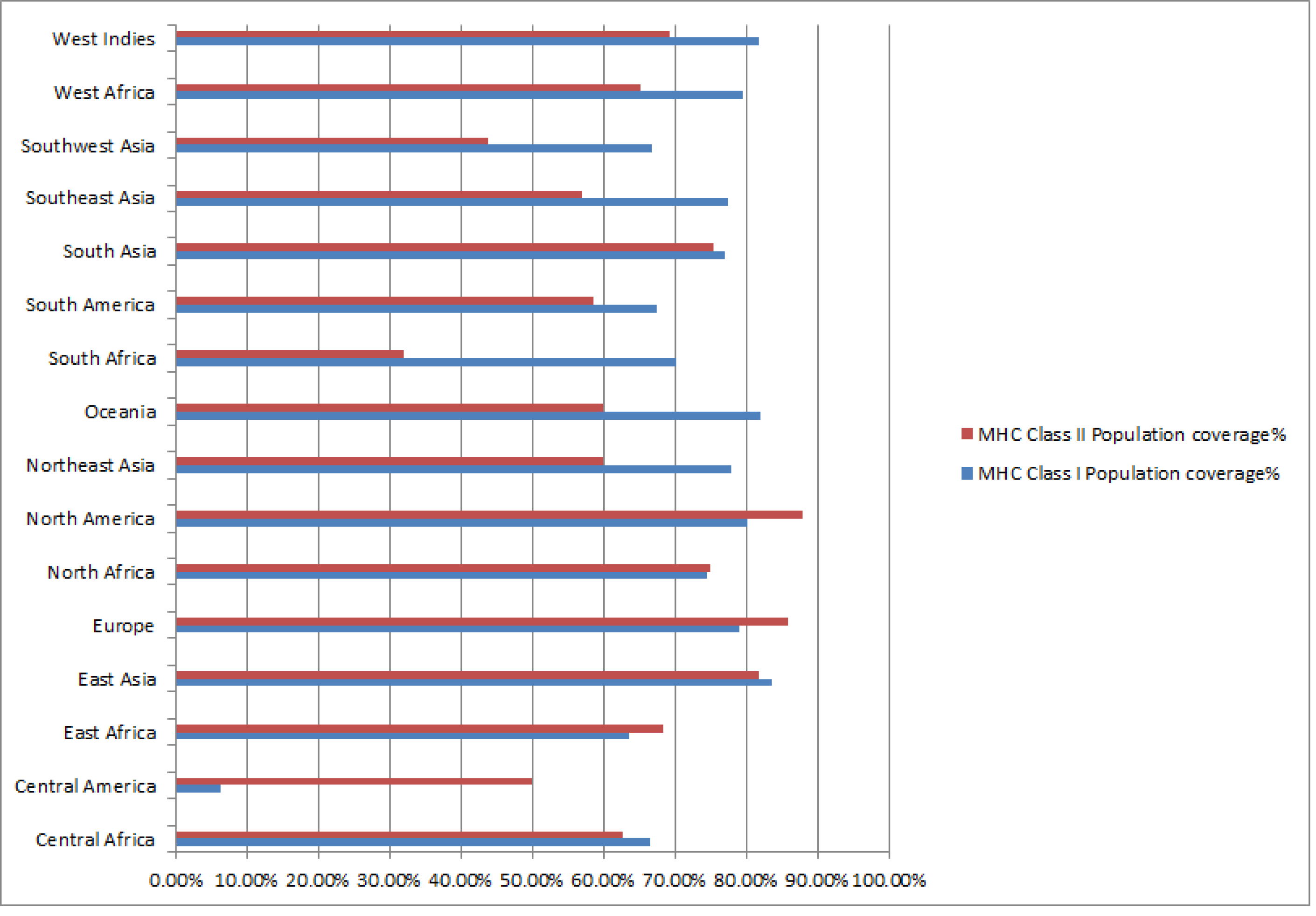
Population coverage of MHC class I & II epitopes in selected geographical regions.

### 5. Structural model and physicochemical properties of Multi Epitope Subunit Vaccine (MESV)

The constructed Multi Epitope Subunit Vaccine (MESV) comprises of 280 amino acids from 15 antigenic B-cell and T-cell epitopes that have been linked with an immuno adjuvant. The tertiary structure of the multiple epitope vaccine was constructed using I-TASSER server with a C-score of -4.11 and estimated TM-score of 0.20 ± 0.05 and was further structurally validated using Ramachandran statistics and ProSA-web server. The Ramachandran statistics of 3D model was 66.5% in core, 26.4% in allowed, 5.0% in generously allowed and 2.1% in disallowed region. After rounds of energy refinement, the Ramachandran statistics were improved to 82.6% in core, 13.6% in allowed, 0.8% in generously allowed and 2.9% in disallowed region (Figure 6). The MESV structural model had a Z-score of -2.32, indicating that it was a rather acceptable model. The physiochemical attributes of MESV are as follow: No. of amino acids (280), Mol. Wt. 31277.77 Da, pI 10.19, estimated half-life 30 hours in eukaryotes, instability index 36.51, aliphatic index 71.61 and GRAVY score of -0.139.

**Figure 6:**
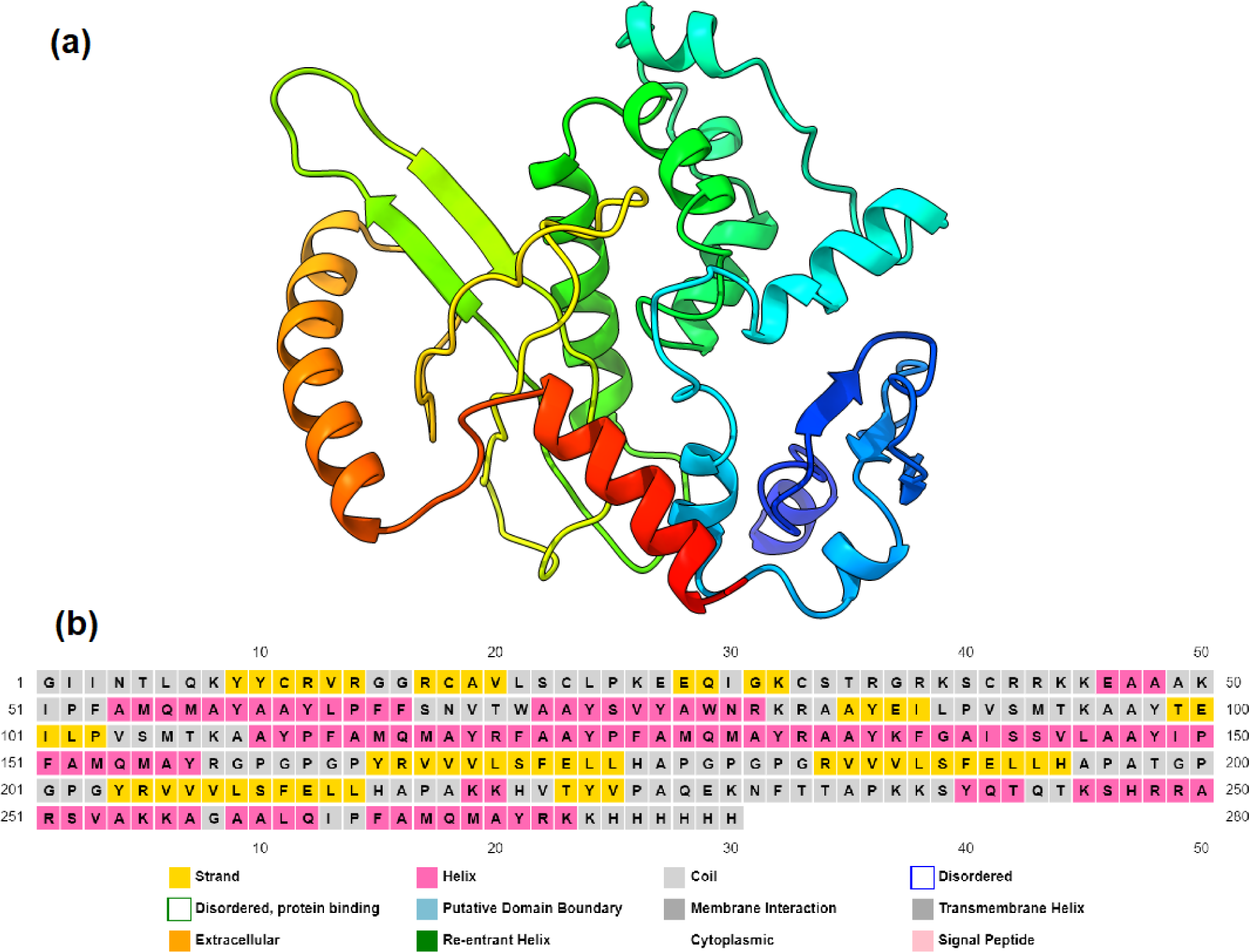
(a) 3D structure model of MESV shown in cartoon representation, and (b) Secondary structure prediction of the MESV.

### 6. Molecular Docking between the Vaccine and the toll-like receptor (TLR5)

TLR5 was chosen for its immunomodulatory potential to induce IFN-gamma. This wa demonstrated in the study, where our chosen CD4+ epitopes induced both Th1 and Th2 cytokines. The vaccine’s molecular interaction with TLR5 (PDB: 3J0A) was studied, taking into account their refined binding energies and numerous interacting residues. The possible orientation and interactions between the MESV and TLR5 are shown in Figure 7a and 7b respectively. The predicted binding free energy of complex was -48.61 (kcal/mol).

**Figure 7:**
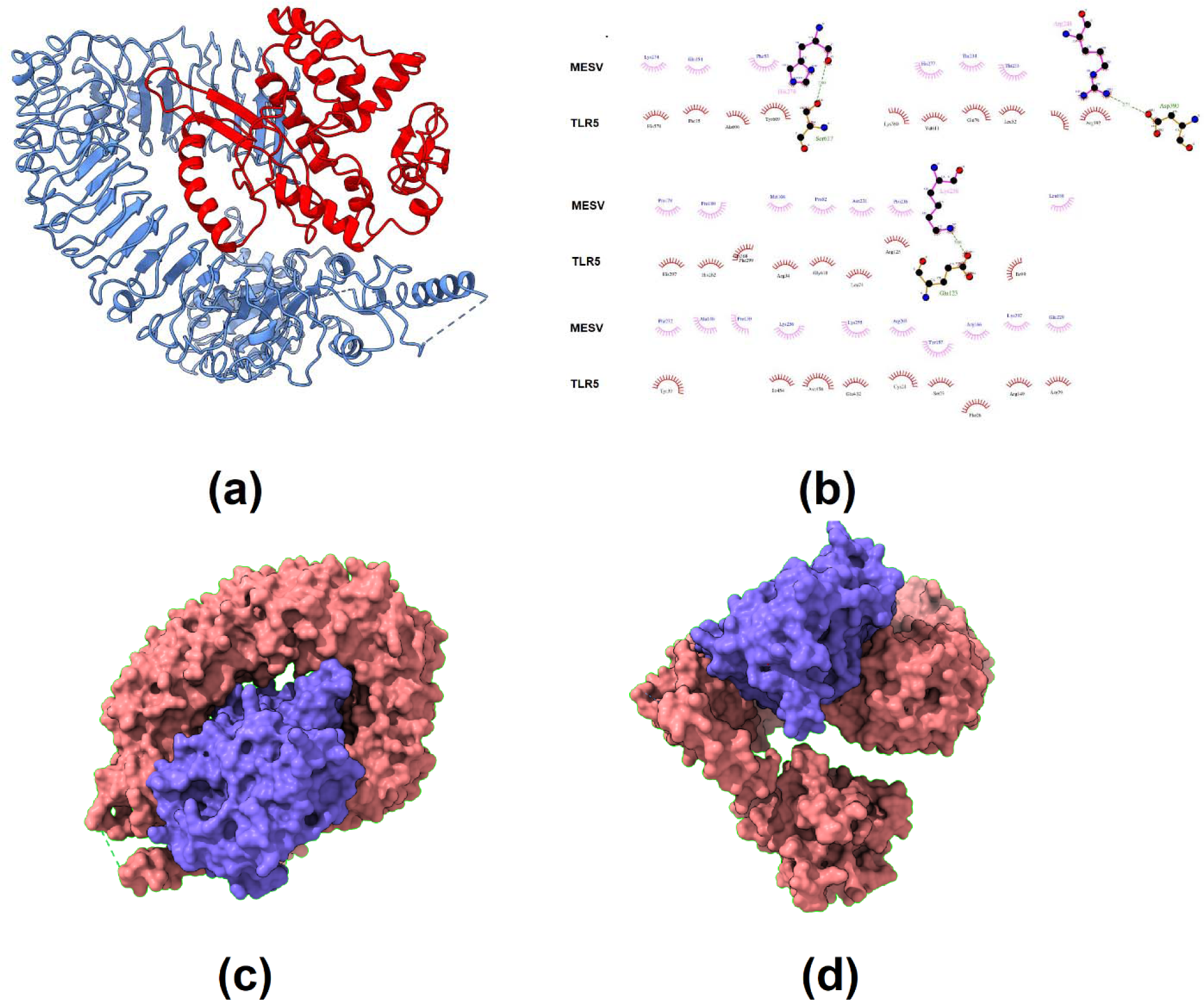
(a) Docked complex of MESV (red) and TLR5 receptor (blue); (b) Interactions of MESV and TLR5 receptor (c, d) Docked complex of MESV (blue) and TLR5 receptor (red) in surface representation at different angles.

### 7. In-silico codon optimization and cloning

The optimal vaccine codon sequence was 840 nucleotides long. The GC content of the cDNA sequence and codon adaptive index was calculated to be 52.14 percent, which is still within th recommended range of 30–70% for effective translational efficiency. The computed codon adaptability index was 1, which is within the range of 0.8–1.0, indicating that the vaccine designs were effectively expressed in *E. coli*. The optimized nucleotide sequence was cloned into the pET28a (+) vector with *Nde*I and *Eag*I restriction enzymes (Supplementary figure S2).

### 8. MD simulation and Immune simulation of the MESV

The radius of gyration shows the rigidity of the vaccine and the Rg value of the vaccine-TLR5 construct was 36.275. The eigenvalue of the vaccine-TLR5 complex was 1.697699e-06, value is low which shows the easier deformation of the complex, which means that the vaccine-TLR5 construct will activate the immune cascade. At some points, the B factor value of the complex is high, which directly corresponds to the higher deformability. The MD simulations results are shown in Figure 8. The immune simulation of the vaccine was also carried out for simulating the immune response post vaccination. At the time of the administration of every dose of the MESV peptide vaccine, there is a sharp increase in the antibody response and a co-currently decrease in the antigen level. The antibody response was significantly higher i.e. IgM > IgG humoral response highlighting the serocon-version (Figure 9A). Also, following the second dose, IFN-γ response was higher (accompanying with both CD8+ T-cell and CD4+ Th 1 response), and also IL-10 & TGF-b cytokines response related with T-reg phenotype (Figure 9B). Overall, the immune simulations results showed with each dose revealed that there was simultaneous increase in immune response. B-cell population per cell analysis revealed that the overall B-cell population and B-cell memory responses were higher and stable showing minimal decay for over 300 days (Figure 9C). After the second dose, there was a simultaneous rise in Th effector cell phenotype, and lower Th-memory readout (Figure 9D). Along with these, a concomitant increase in natural killer cells activities (Figure 9E) throughout the simulation. To summarize the results, the points mentioned above is a good performance indicator of the vaccine construct showing its capability to stimulate the correct immunological compartment for an effective response.

**Figure 8:**
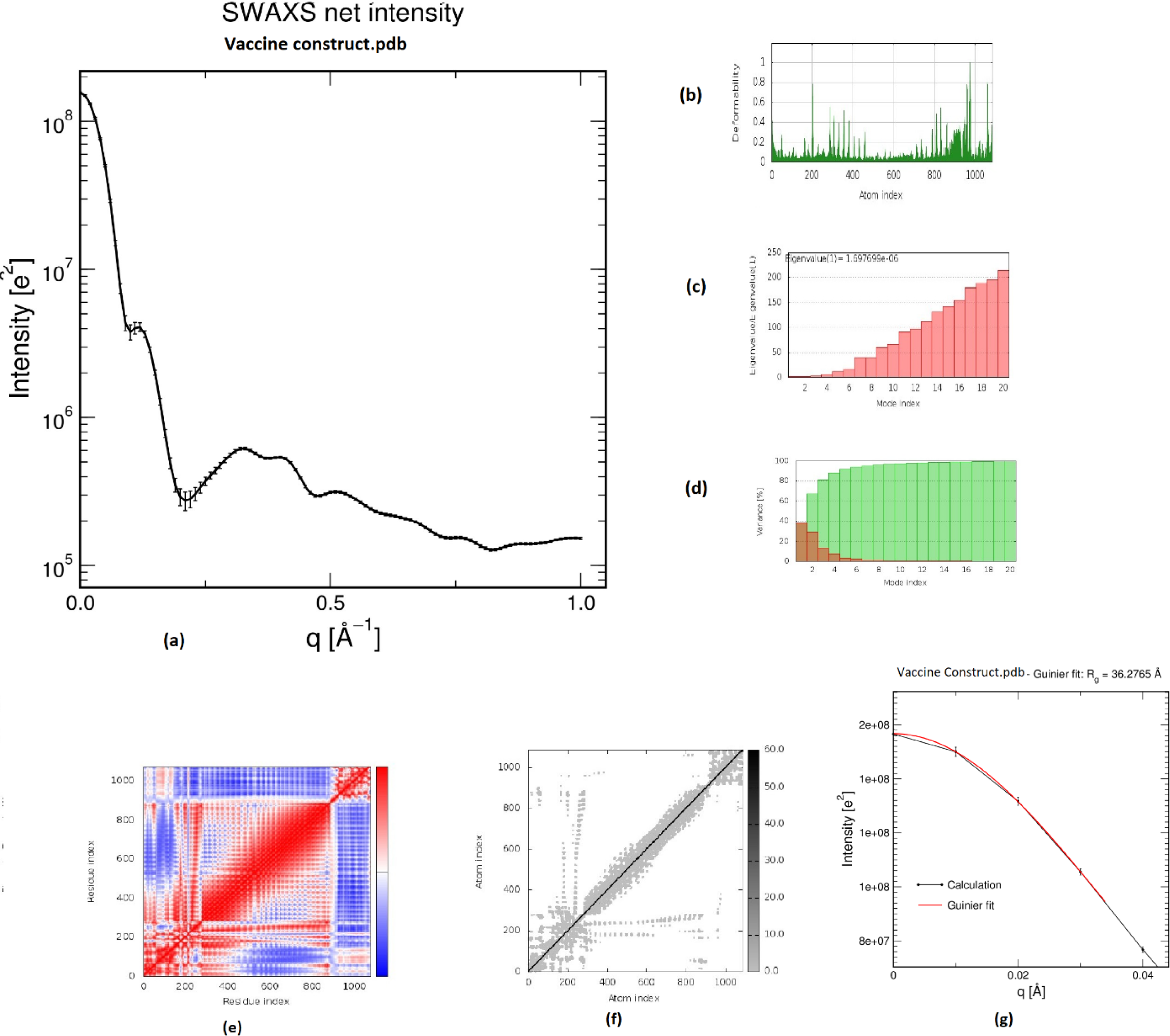
The MD simulation results of the MESV vaccine. (a) SAWXS net intensity of MSEV-TLR5 complex; molecular dynamics simulation of the MESV-TLR5 complex, showing (b) deformability; (c) eigenvalue; and (d) variance; (e) Covariance matrix indicates coupling between pairs of residues (red), uncorrelated (white) or anti-correlated (blue) motions; (f) elastic network analysis which defines which pairs of atoms are connected by springs; and (g) radius of gyration plot showing the calculated values vs. Guinier fit.

**Figure 9:**
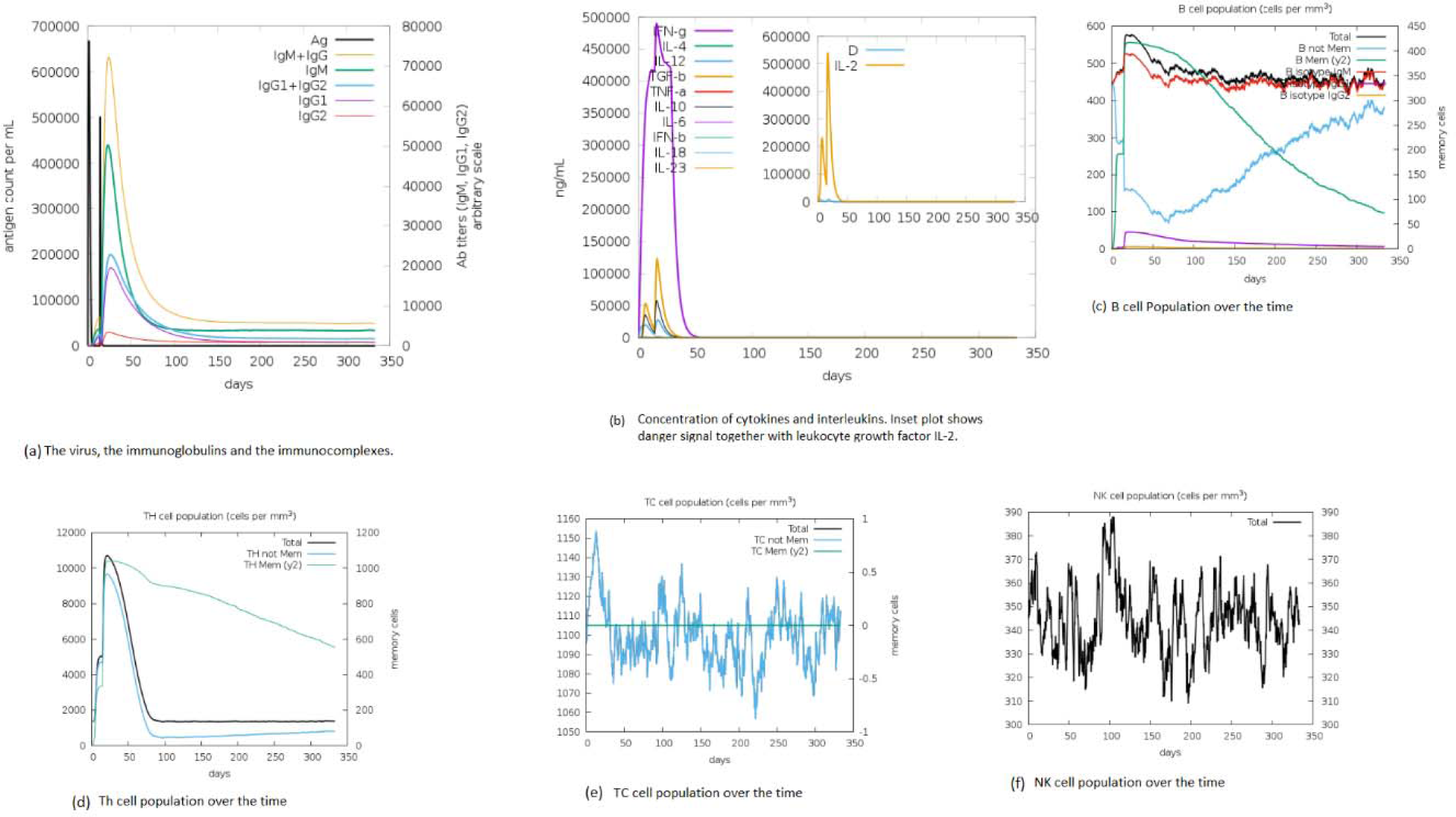
Results of MESV immune simulations. (a) plot of antigen level along with immunoglobin profile; (b) induced array of cytokines; (c) B-cell population over the time (d) Th cell population over the time (e) Tc cell population over the time (f) Natural Killer cells (total count).

## Acknowledgements

The authors acknowledge the Gujarat State Biotechnology Mission (GSBTM), Department of Science & Technology (DST), Gujarat for financial support and resources. The authors also acknowledge the Institute of Advanced Research, Gandhinagar for resources and infrastructure facilities. The authors are also grateful to the Gujarat Council on Science and Technology (GUJCOST), Department of Science & Technology (DST), Gujarat for the PARAM SHAVAK supercomputing facility and DST - Innovation, Technology Development and Deployment scheme grant DST/TMD(EWO)OWUIS-2018/RS-20(G) for resource support.

## Notes

### Competing Interest Statement

The authors have declared no competing interest.

### Summary of Updates

A section and introduction has been added and a image for insilico cloning construct has been included in supplementary section.

## References

1. Báez-Santos, Y. M., St. John, S. E., & Mesecar, A. D. (2015). The SARS-coronavirus papain-like protease: Structure, function and inhibition by designed antiviral compounds. In Antiviral Research (Vol. 115, pp. 21–38). Elsevier B.V. https://doi.org/10.1016/j.antiviral.2014.12.015

2. Behmard, E., Soleymani, B., Najafi, A., & Barzegari, E. (2020). Immunoinformatic design of a COVID-19 subunit vaccine using entire structural immunogenic epitopes of SARS-CoV-2. https://doi.org/10.1038/s41598-020-77547-4

3. Chan, J. F. W., Yuan, S., Kok, K. H., To, K. K. W., Chu, H., Yang, J., Xing, F., Liu, J., Yip, C. C. Y., Poon, R. W. S., Tsoi, H. W., Lo, S. K. F., Chan, K. H., Poon, V. K. M., Chan, W. M., Ip, J. D., Cai, J. P., Cheng, V. C. C., Chen, H., … Yuen, K. Y. (2020). A familial cluster of pneumonia associated with the 2019 novel coronavirus indicating person-to-person transmission: a study of a family cluster. The Lancet, 395(10223), 514–523. https://doi.org/10.1016/S0140-6736(20)30154-9

4. Cui, J., Li, F., & Shi, Z. L. (2019). Origin and evolution of pathogenic coronaviruses. Nature Reviews. Microbiology, 17(3), 181–192. https://doi.org/10.1038/S41579-018-0118-9

5. Dai, L., & Gao, G. F. (2020). Viral targets for vaccines against COVID-19. Nature Reviews Immunology 2020 21:2, 21(2), 73–82. https://doi.org/10.1038/s41577-020-00480-0

6. De Wilde, A. H., Snijder, E. J., Kikkert, M., Van Hemert, M. J., De Wilde, A. H., Snijder, Á. E. J., Kikkert, Á. M., & Van Hemert, Á. M. J. (2017). Host Factors in Coronavirus Replication. Current Topics in Microbiology and Immunology, 419, 1–42. https://doi.org/10.1007/82_2017_25

7. Defay, T., & Cohen, F. E. (1995). Evaluation of current techniques for Ab initio protein structure prediction. *Proteins: Structure*, Function, and Genetics, 23(3), 431–445. https://doi.org/10.1002/prot.340230317

8. Duffy, S. (2018). Why are RNA virus mutation rates so damn high? PLOS Biology, 16(8), e3000003. https://doi.org/10.1371/JOURNAL.PBIO.3000003

9. Harvey, W. T., Carabelli, A. M., Jackson, B., Gupta, R. K., Thomson, E. C., Harrison, E. M., Ludden, C., Reeve, R., Rambaut, A., Peacock, S. J., & Robertson, D. L. (2021). SARS-CoV-2 variants, spike mutations and immune escape. Nature Reviews Microbiology 2021 19:7, 19(7), 409–424. https://doi.org/10.1038/s41579-021-00573-0

10. Hoffmann, M., Kleine-Weber, H., Schroeder, S., Krüger, N., Herrler, T., Erichsen, S., Schiergens, T. S., Herrler, G., Wu, N. H., Nitsche, A., Müller, M. A., Drosten, C., & Pöhlmann, S. (2020). SARS-CoV-2 Cell Entry Depends on ACE2 and TMPRSS2 and Is Blocked by a Clinically Proven Protease Inhibitor. Cell, 181(2). https://doi.org/10.1016/j.cell.2020.02.052

11. Hui, D. S., I Azhar, E., Madani, T. A., Ntoumi, F., Kock, R., Dar, O., Ippolito, G., Mchugh, T. D., Memish, Z. A., Drosten, C., Zumla, A., & Petersen, E. (2020). The continuing 2019-nCoV epidemic threat of novel coronaviruses to global health — The latest 2019 novel coronavirus outbreak in Wuhan, China. In International Journal of Infectious Diseases (Vol. 91, pp. 264–266). Elsevier B.V. https://doi.org/10.1016/j.ijid.2020.01.009

12. Hwang, S. S., Lim, J., Yu, Z., Kong, P., Sefik, E., Xu, H., Harman, C. C. D., Kim, L. K., Lee, G. R., Li, H. B., & Flavell, R. A. (2020). Cryo-EM structure of the 2019-nCoV spike in the prefusion conformation. *Science (New York*, N.Y.), 367(6483), 1255–1260. https://doi.org/10.1126/SCIENCE.ABB2507

13. Ishack, S., & Lipner, S. R. (2021). Bioinformatics and immunoinformatics to support COVID-19 vaccine development. Journal of Medical Virology, 93(9), 5209–5211. https://doi.org/10.1002/JMV.27017/FORMAT/PDF

14. Jia, Z., & Gong, W. (2021). Will Mutations in the Spike Protein of SARS-CoV-2 Lead to the Failure of COVID-19 Vaccines? Journal of Korean Medical Science, 36(18), 1–11. https://doi.org/10.3346/JKMS.2021.36.E124

15. Lauring, A. S., & Hodcroft, E. B. (2021). Genetic Variants of SARS-CoV-2—What Do They Mean? JAMA, 325(6), 529–531. https://doi.org/10.1001/JAMA.2020.27124

16. Liu, L., Wang, P., Nair, M. S., Yu, J., Rapp, M., Wang, Q., Luo, Y., Chan, J. F. W., Sahi, V., Figueroa, A., Guo, X. V., Cerutti, G., Bimela, J., Gorman, J., Zhou, T., Chen, Z., Yuen, K. Y., Kwong, P. D., Sodroski, J. G., … Ho, D. D. (2020). Potent neutralizing antibodies against multiple epitopes on SARS-CoV-2 spike. Nature 2020 584:7821, 584(7821), 450–456. https://doi.org/10.1038/s41586-020-2571-7

17. Lu, R., Zhao, X., Li, J., Niu, P., Yang, B., Wu, H., Wang, W., Song, H., Huang, B., Zhu, N., Bi, Y., Ma, X., Zhan, F., Wang, L., Hu, T., Zhou, H., Hu, Z., Zhou, W., Zhao, L., … Tan, W. (2020). Genomic characterisation and epidemiology of 2019 novel coronavirus: implications for virus origins and receptor binding. The Lancet, 395, 565– 574. https://doi.org/10.1016/S0140-6736(20)30251-8

18. Meredith, L. W., Hamilton, W. L., Warne, B., Houldcroft, C. J., Hosmillo, M., Jahun, A. S., Curran, M. D., Parmar, S., Caller, L. G., Caddy, S. L., Khokhar, F. A., Yakovleva, A., Hall, G., Feltwell, T., Forrest, S., Sridhar, S., Weekes, M. P., Baker, S., Brown, N., … Goodfellow, I. (2020). Rapid implementation of SARS-CoV-2 sequencing to investigate cases of health-care associated COVID-19: a prospective genomic surveillance study. The Lancet. Infectious Diseases, 20(11), 1263–1272. https://doi.org/10.1016/S1473-3099(20)30562-4

19. Naqvi, A. A. T., Fatima, K., Mohammad, T., Fatima, U., Singh, I. K., Singh, A., Atif, S. M., Hariprasad, G., Hasan, G. M., & Hassan, M. I. (2020). Insights into SARS-CoV-2 genome, structure, evolution, pathogenesis and therapies: Structural genomics approach. In Biochimica et Biophysica Acta - Molecular Basis of Disease (Vol. 1866, Issue 10, p. 165878). Elsevier B.V. https://doi.org/10.1016/j.bbadis.2020.165878

20. Nicola, M., Alsafi, Z., Sohrabi, C., Kerwan, A., Al-Jabir, A., Iosifidis, C., Agha, M., & Agha, R. (2020). The socio-economic implications of the coronavirus pandemic (COVID-19): A review. *International Journal of Surgery (London*, England), 78, 185. https://doi.org/10.1016/J.IJSU.2020.04.018

21. Piccoli, L., Park, Y. J., Tortorici, M. A., Czudnochowski, N., Walls, A. C., Beltramello, M., Silacci-Fregni, C., Pinto, D., Rosen, L. E., Bowen, J. E., Acton, O. J., Jaconi, S., Guarino, B., Minola, A., Zatta, F., Sprugasci, N., Bassi, J., Peter, A., De Marco, A., … Veesler, D. (2020). Mapping Neutralizing and Immunodominant Sites on the SARS-CoV-2 Spike Receptor-Binding Domain by Structure-Guided High-Resolution Serology. Cell, 183(4), 1024–1042.e21. https://doi.org/10.1016/J.CELL.2020.09.037

22. Rees-Spear, C., Muir, L., Griffith, S. A., Heaney, J., Aldon, Y., Snitselaar, J. L., Thomas, P., Graham, C., Seow, J., Lee, N., Rosa, A., Roustan, C., Houlihan, C. F., Sanders, R. W., Gupta, R. K., Cherepanov, P., Stauss, H. J., Nastouli, E., Doores, K. J., … McCoy, L. E. (2021). The effect of spike mutations on SARS-CoV-2 neutralization. Cell Reports, 34(12). https://doi.org/10.1016/J.CELREP.2021.108890

23. Rota, P. A., Oberste, M. S., Monroe, S. S., Nix, W. A., Campagnoli, R., Icenogle, J. P., Peñaranda, S., Bankamp, B., Maher, K., Chen, Mhsin, Tong, S., Tamin, A., Lowe, L., Frace, M., DeRisi, J. L., Chen, Q., Wang, D., Erdman, D. D., Peret, T. C. T., … Bellini, W. J. (2003). Characterization of a novel coronavirus associated with severe acute respiratory syndrome. Science, 300(5624), 1394–1399. https://doi.org/10.1126/science.1085952

24. Saha, R. P., Sharma, A. R., Singh, M. K., Samanta, S., Bhakta, S., Mandal, S., Bhattacharya, M., Lee, S. S., & Chakraborty, C. (2020). Repurposing Drugs, Ongoing Vaccine, and New Therapeutic Development Initiatives Against COVID-19. Frontiers in Pharmacology, 11, 1258. https://doi.org/10.3389/FPHAR.2020.01258/BIBTEX

25. SARS-CoV-2 Variant Classifications and Definitions. (2021). https://www.cdc.gov/coronavirus/2019-ncov/variants/variant-info.html

26. Science Brief: Omicron (B.1.1.529) Variant | CDC. (2021). https://www.cdc.gov/coronavirus/2019-ncov/science/science-briefs/scientific-brief-omicron-variant.html

27. Socio-economic impact of COVID-19 | United Nations Development Programme. (2020). UNDP. https://www.undp.org/coronavirus/socio-economic-impact-covid-19

28. Toyoshima, Y., Nemoto, K., Matsumoto, S., Nakamura, Y., & Kiyotani, K. (2020). SARS-CoV-2 genomic variations associated with mortality rate of COVID-19. Journal of Human Genetics 2020 65:12, 65(12), 1075–1082. https://doi.org/10.1038/s10038-020-0808-9

29. Tracking SARS-CoV-2 variants. (2021). https://www.who.int/en/activities/tracking-SARS-CoV-2-variants/

30. Wahid, M., Jawed, A., Mandal, R. K., Dailah, H. G., Janahi, E. M., Dhama, K., Somvanshi, P., & Haque, S. (2021). Variants of SARS-CoV-2, their effects on infection, transmission and neutralization by vaccine induced antibodies. European Review for Medical and Pharmacological Sciences, 25(18), 5857–5864. https://doi.org/10.26355/EURREV_202109_26805

31. Walls, A. C., Park, Y. J., Tortorici, M. A., Wall, A., McGuire, A. T., & Veesler, D. (2020). Structure, Function, and Antigenicity of the SARS-CoV-2 Spike Glycoprotein. Cell, 181(2), 281-292.e6. https://doi.org/10.1016/J.CELL.2020.02.058

32. Wan, Y., Shang, J., Graham, R., Baric, R. S., & Li, F. (2020). Receptor Recognition by the Novel Coronavirus from Wuhan: an Analysis Based on Decade-Long Structural Studies of SARS Coronavirus. Journal of Virology, 94(7). https://doi.org/10.1128/JVI.00127-20

33. Wang, C., Horby, P. W., Hayden, F. G., & Gao, G. F. (2020). A novel coronavirus outbreak of global health concern. In The Lancet (Vol. 395, Issue 10223, pp. 470–473). Lancet Publishing Group. https://doi.org/10.1016/S0140-6736(20)30185-9

34. Wu, F., Zhao, S., Yu, B., Chen, Y. M., Wang, W., Song, Z. G., Hu, Y., Tao, Z. W., Tian, J. H., Pei, Y. Y., Yuan, M. L., Zhang, Y. L., Dai, F. H., Liu, Y., Wang, Q. M., Zheng, J. J., Xu, L., Holmes, E. C., & Zhang, Y. Z. (2020). A new coronavirus associated with human respiratory disease in China. Nature, 579(7798), 265–269. https://doi.org/10.1038/s41586-020-2008-3

35. Xu, H., Zhong, L., Deng, J., Peng, J., Dan, H., Zeng, X., Li, T., & Chen, Q. (2020). High expression of ACE2 receptor of 2019-nCoV on the epithelial cells of oral mucosa. International Journal of Oral Science 2020 12:1, 12(1), 1–5. https://doi.org/10.1038/s41368-020-0074-x

36. Yuan, M., Wu, N. C., Zhu, X., Lee, C. C. D., So, R. T. Y., Lv, H., Mok, C. K. P., & Wilson, I. A. (2020). A highly conserved cryptic epitope in the receptor binding domains of SARS-CoV-2 and SARS-CoV. *Science (New York*, N.Y.), 368(6491), 630–633. https://doi.org/10.1126/SCIENCE.ABB7269

37. Zhou, P., Yang, X.-L., Wang, X.-G., Hu, B., Zhang, L., Zhang, W., Si, H.-R., Zhu, Y., Li, B., Huang, C.-L., Chen, H.-D., Chen, J., Luo, Y., Guo, H., Jiang, R.-D., Liu, M.-Q., Chen, Y., Shen, X.-R., Wang, X., … Shi, Z.-L. (2020). Discovery of a novel coronavirus associated with the recent pneumonia outbreak in humans and its potential bat origin. BioRxiv, 2020.01.22.914952. https://doi.org/10.1101/2020.01.22.914952

38. Zhu, N., Zhang, D., Wang, W., Li, X., Yang, B., Song, J., Zhao, X., Huang, B., Shi, W., Lu, R., Niu, P., Zhan, F., Ma, X., Wang, D., Xu, W., Wu, G., Gao, G. F., & Tan, W. (2020). A Novel Coronavirus from Patients with Pneumonia in China, 2019. The New England Journal of Medicine, 382(8), 727–733. https://doi.org/10.1056/NEJMOA2001017

39. Ziebuhr, J., Snijder, E. J., & Gorbalenya, A. E. (2000). Virus-encoded proteinases and proteolytic processing in the Nidovirales. In Journal of General Virology (Vol. 81, Issue 4, pp. 853–879). Society for General Microbiology. https://doi.org/10.1099/0022-1317-81-4-853

