## Supplemental Figures for "Designing multi-epitope based peptide vaccine targeting spike protein SARS-CoV-2 B1.1.529 (Omicron) variant using computational approaches"

```
B.1.1.529_Omicron    η1  TT  β1  β2
                     1  10  20  30  40  50  60
B.1.1.529_Omicron    MFVFLVLLPLVSSQCVNLTTRTQLPPAYTNSFTRGVYYPDKVFRSSVLHSTQDLFLPFFS
NC_045512.2_Wuhan    MFVFLVLLPLVSSQCVNLTTRTQLPPAYTNSFTRGVYYPDKVFRSSVLHSTQDLFLPFFS
B.1.617.2_Delta      MFVFLVLLPLVSSQCVNLRTRTQLPPAYTNSFTRGVYYPDKVFRSSVLHSTQDLFLPFFS
                     1

B.1.1.529_Omicron    β3  β4  β5  β6  TT  β7
                     70  80  90  100  110
B.1.1.529_Omicron    NVTWFHVI..SGTNGTKRFDNPVLPFNDGVYFASIEKSNIIRGWIFGTTLDSKTQSLLIV
NC_045512.2_Wuhan    NVTWFHAIHVSGTNGTKRFDNPVLPFNDGVYFASTEKSNIIRGWIFGTTLDSKTQSLLIV
B.1.617.2_Delta      NVTWFHAIHVSGTNGTKRFDNPVLPFNDGVYFASTEKSNIIRGWIFGTTLDSKTQSLLIV

B.1.1.529_Omicron    β8  TT  TT  β9
                     120  130  140  150  160  170
B.1.1.529_Omicron    NNATNVVIKVCEFQFCNDPFLD...HKNNKSWMESEFRVYSSANNCTFEYVSQPFLMDLE
NC_045512.2_Wuhan    NNATNVVIKVCEFQFCNDPFLGVYYHKNNKSWMESEFRVYSSANNCTFEYVSQPFLMDLE
B.1.617.2_Delta      NNATNVVIKVCEFQFCNDPFLXVYYHKNNKSWMESEFRVYSSANNCTFEYVSQPFLMDLE
                     2  1  2

B.1.1.529_Omicron    α1  β10  TT  β11  β12
                     180  190  200  210  220  230
B.1.1.529_Omicron    GKQGNFKNLREFVFKNIDGYFKIYSKHTPI.IVREPEDLPQGFSALEPLVDLPIGINITR
NC_045512.2_Wuhan    GKQGNFKNLREFVFKNIDGYFKIYSKHTPINLVR...DLPQGFSALEPLVDLPIGINITR
B.1.617.2_Delta      GKQGNFKNLREFVFKNIDGYFKIYSKHTPINLVR...DLPQGFSALEPLVDLPIGINITR

B.1.1.529_Omicron    β13  TTT  β14  β15  TT  β16  TT
                     240  250  260  270  280  290
B.1.1.529_Omicron    FQTLLALHRSYLTPGDSSSGWTAGAAAYYVGYLQPRTFLLKYNENGTITDAVDCALDPLS
NC_045512.2_Wuhan    FQTLLALHRSYLTPGDSSSGWTAGAAAYYVGYLQPRTFLLKYNENGTITDAVDCALDPLS
B.1.617.2_Delta      FQTLLALHRSYLTPGDSSSGWTAGAAAYYVGYLQPRTFLLKYNENGTITDAVDCALDPLS
                     3

B.1.1.529_Omicron    α2  β17  β18  α3  η2  β19
                     300  310  320  330  340  350
B.1.1.529_Omicron    ETKCTLKSFTVEKGIYQTSNFRVQPTESIVRFPNITNLCPFDEVFNATRFASVYAWNRKR
NC_045512.2_Wuhan    ETKCTLKSFTVEKGIYQTSNFRVQPTESIVRFPNITNLCPFGEVFNATRFASVYAWNRKR
B.1.617.2_Delta      ETKCTLKSFTVEKGIYQTSNFRVQPTESIVRFPNITNLCPFGEVFNATRFASVYAWNRKR
                     3  4

B.1.1.529_Omicron    β20  α4  β21  η3  β22  β23  α5  TT
                     360  370  380  390  400  410
B.1.1.529_Omicron    ISNCVADYSVLYNLAPFFTFKCYGVSPTKLNDLCFTNVYADSFVIRGDEVRQIAPGQTGN
NC_045512.2_Wuhan    ISNCVADYSVLYNSASFSTFKCYGVSPTKLNDLCFTNVYADSFVIRGDEVRQIAPGQTGK
B.1.617.2_Delta      ISNCVADYSVLYNSASFSTFKCYGVSPTKLNDLCFTNVYADSFVIRGDEVRQIAPGQTGK
                     4  5  6

B.1.1.529_Omicron    α6  TT  β24  α7  TT  β25  TT  β26
                     420  430  440  450  460  470
B.1.1.529_Omicron    IADYNYKLPDDFTGCVIAWNSNKLDSKVSGNYNYLYRLFRKSNLKPFERDISTEIYQAGN
NC_045512.2_Wuhan    IADYNYKLPDDFTGCVIAWNSNNLDSKVGGNYNYLYRLFRKSNLKPFERDISTEIYQAGS
B.1.617.2_Delta      IADYNYKLPDDFTGCVIAWNSNNLDSKVGGNYNYRYRLFRKSNLKPFERDISTQIYQAGS
                     5

B.1.1.529_Omicron    TT  β27  β28  TT  TT  β29  β30
                     480  490  500  510  520  530
B.1.1.529_Omicron    KPCNGVAGFNCYFPLKSYSFRPTYGVGHQPYRVVVLSFELLHAPATVCGPKKSTNLVKNK
NC_045512.2_Wuhan    TPCNGVEGFNCYFPLQSYGFQPTNGVGYQPYRVVVLSFELLHAPATVCGPKKSTNLVKNK
B.1.617.2_Delta      KPCNGVEGFNCYFPLQSYGFQPTNGVGYQPYRVVVLSFELLHAPATVCGPKKSTNLVKNK
                     7  7  6

B.1.1.529_Omicron    β31  TT  β32  TT  β33  TT  β34  TTT  β35  β36
                     540  550  560  570  580  590
B.1.1.529_Omicron    CVNFNFNGLKGTGVLTESNKKFLPFQQFGRDIADTTDAVRDPQTLEILDITPCSFGGVSV
NC_045512.2_Wuhan    CVNFNFNGLTGTGVLTESNKKFLPFQQFGRDIADTTDAVRDPQTLEILDITPCSFGGVSV
B.1.617.2_Delta      CVNFNFNGLTGTGVLTESNKKFLPFQQFGRDIADTTDAVRDPQTLEILDITPCSFGGVSV
                     8  8

B.1.1.529_Omicron    β37  η4  α8  β38  TT  β39  β40
                     600  610  620  630  640  650
B.1.1.529_Omicron    ITPGTNTSNQVAVLYQGVNCTEVPVAIHADQLTPTWRVYSTGSNVFQTRAGCLIGAEYVN
NC_045512.2_Wuhan    ITPGTNTSNQVAVLYQDVNCTEVPVAIHADQLTPTWRVYSTGSNVFQTRAGCLIGAEHVN
B.1.617.2_Delta      ITPGTNTSNQVAVLYQGVNCTEVPVAIHADQLTPTWRVYSTGSNVFQTRAGCLIGAEHVN
                     9  9
```

```
B.1.1.529_Omicron              β41 TT β42                          β43                        TT β44
                               660         670         680         690         700         710
B.1.1.529_Omicron   NSYECDIPIGAGICASYQTQTKSHRRARSVASQSIIAYTMSLGAENSVAYSNNSIAIPTN
NC_045512.2_Wuhan   NSYECDIPIGAGICASYQTQTNSPRRARSVASQSIIAYTMSLGAENSVAYSNNSIAIPTN
B.1.617.2_Delta     NSYECDIPIGAGICASYQTQTNSRRRARSVASQSIIAYTMSLGAENSVAYSNNSIAIPTN
                         0         0

B.1.1.529_Omicron   β45                 β46  α9        α10              α11
                          720         730         740         750         760         770
B.1.1.529_Omicron   FTISVTTEILPVSMTKTSVDCTMYICGDSTECSNLLLQYGSFCTQLKRALTGIAVEQDKN
NC_045512.2_Wuhan   FTISVTTEILPVSMTKTSVDCTMYICGDSTECSNLLLQYGSFCTQLNRALTGIAVEQDKN
B.1.617.2_Delta     FTISVTTEILPVSMTKTSVDCTMYICGDSTECSNLLLQYGSFCTQLNRALTGIAVEQDKN
                                                   1     2      2       1

B.1.1.529_Omicron                    TT η5      TT     α12
                          780         790         800         810         820         830
B.1.1.529_Omicron   TQEVFAQVKQIYKTPPIKYFGGFNFSQILPDPSKPSKRSFIEDLLFNKVTLADAGFIKQY
NC_045512.2_Wuhan   TQEVFAQVKQIYKTPPIKDFGGFNFSQILPDPSKPSKRSFIEDLLFNKVTLADAGFIKQY
B.1.617.2_Delta     TQEVFAQVKQIYKTPPIKDFGGFNFSQILPDPSKPSKRSFIEDLLFNKVTLADAGFIKQY

B.1.1.529_Omicron   η6   η7   α13     β47          α14              η8
                          840         850         860         870         880         890
B.1.1.529_Omicron   GDCLGDIAARDLICAQKFKGLTVLPPLLTDEMIAQYTSALLAGTITSGWTFGAGAALQIP
NC_045512.2_Wuhan   GDCLGDIAARDLICAQKFNGLTVLPPLLTDEMIAQYTSALLAGTITSGWTFGAGAALQIP
B.1.617.2_Delta     GDCLGDIAARDLICAQKFNGLTVLPPLLTDEMIAQYTSALLAGTITSGWTFGAGAALQIP
                       3           3

B.1.1.529_Omicron        α15                     α16                    η9      α17
                          900         910         920         930         940         950
B.1.1.529_Omicron   FAMQMAYRFNGIGVTQNVLYENQKLIANQFNSAIGKIQDSLSSTASALGKLQDVVNHNAQ
NC_045512.2_Wuhan   FAMQMAYRFNGIGVTQNVLYENQKLIANQFNSAIGKIQDSLSSTASALGKLQDVVNQNAQ
B.1.617.2_Delta     FAMQMAYRFNGIGVTQNVLYENQKLIANQFNSAIGKIQDSLSSTASALGKLQNVVNQNAQ

B.1.1.529_Omicron                TT    α18                          α19
                          960         970         980         990        1000        1010
B.1.1.529_Omicron   ALNTLVKQLSSKFGAISSVLNDIFSRLDKVEAEVQIDRLITGRLQSLQTYVTQQLIRAAE
NC_045512.2_Wuhan   ALNTLVKQLSSNFGAISSVLNDILSRLDKVEAEVQIDRLITGRLQSLQTYVTQQLIRAAE
B.1.617.2_Delta     ALNTLVKQLSSNFGAISSVLNDILSRLDKVEAEVQIDRLITGRLQSLQTYVTQQLIRAAE

B.1.1.529_Omicron                                        β48     TT  β49          β50
                         1020        1030        1040        1050        1060        1070
B.1.1.529_Omicron   IRASANLAATKMSECVLGQSKRVDFCGKGYHLMSFPQSAPHGVVFLHVTYVPAQEKNFTT
NC_045512.2_Wuhan   IRASANLAATKMSECVLGQSKRVDFCGKGYHLMSFPQSAPHGVVFLHVTYVPAQEKNFTT
B.1.617.2_Delta     IRASANLAATKMSECVLGQSKRVDFCGKGYHLMSFPQSAPHGVVFLHVTYVPAQEKNFTT
                                 4           4

B.1.1.529_Omicron     β51 TT β52    β53    β54        β55    β56    TT TT β57
                         1080        1090        1100        1110        1120        1130
B.1.1.529_Omicron   APAICHDGKAHFPREGVFVSNGTHWFVTQRNFYEPQIITTDNTFVSGNCDVVIGIVNNTV
NC_045512.2_Wuhan   APAICHDGKAHFPREGVFVSNGTHWFVTQRNFYEPQIITTDNTFVSGNCDVVIGIVNNTV
B.1.617.2_Delta     APAICHDGKAHFPREGVFVSNGTHWFVTQRNFYEPQIITTDNTFVSGNCDVVIGIVNNTV
                         5                                             5

B.1.1.529_Omicron             α20            α21                   α22
                         1140        1150        1160        1170        1180        1190
B.1.1.529_Omicron   YDPLQPELDSFKEELDKYFKNHTSPDVDLGDISGINASVVNIQKEIDRLNEVAKNLNESL
NC_045512.2_Wuhan   YDPLQPELDSFKEELDKYFKNHTSPDVDLGDISGINASVVNIQKEIDRLNEVAKNLNESL
B.1.617.2_Delta     YDPLQPELDSFKEELDKYFKNHTSPDVDLGDISGINASVVNIQKEIDRLNEVAKNLNESL

B.1.1.529_Omicron                            α23 β58                  TT
                         1200        1210        1220        1230        1240        1250
B.1.1.529_Omicron   IDLQELGKYEQYIKWPWYIWLGFIAGLIAIVMVTIMLCCMTSCCSCLKGCCSCGSCCKFD
NC_045512.2_Wuhan   IDLQELGKYEQYIKWPWY..........................................
B.1.617.2_Delta     IDLQELGKYEQYIKWPWYIWLGFIAGLIAIVMVTIMLCCMTSCCSCLKGCCSCGSCCKFD

B.1.1.529_Omicron
                         1260        1270
B.1.1.529_Omicron   EDDSEPVLKGVKLHYT
NC_045512.2_Wuhan   ................
B.1.617.2_Delta     EDDSEPVLKGVKLHYT
```

**Figure S1: Multiple structure-based sequence alignment of SARS-CoV-2 spike protein sequences from Wuhan reference strain, Delta variant and Omicron variant. SARS-CoV-2 spike protein from Omicron variant was used as a reference and compared with Wuhan and Delta strain. Domain assignments are marked in the alignment output. The different residues in the alignment are indicated by the white box.**

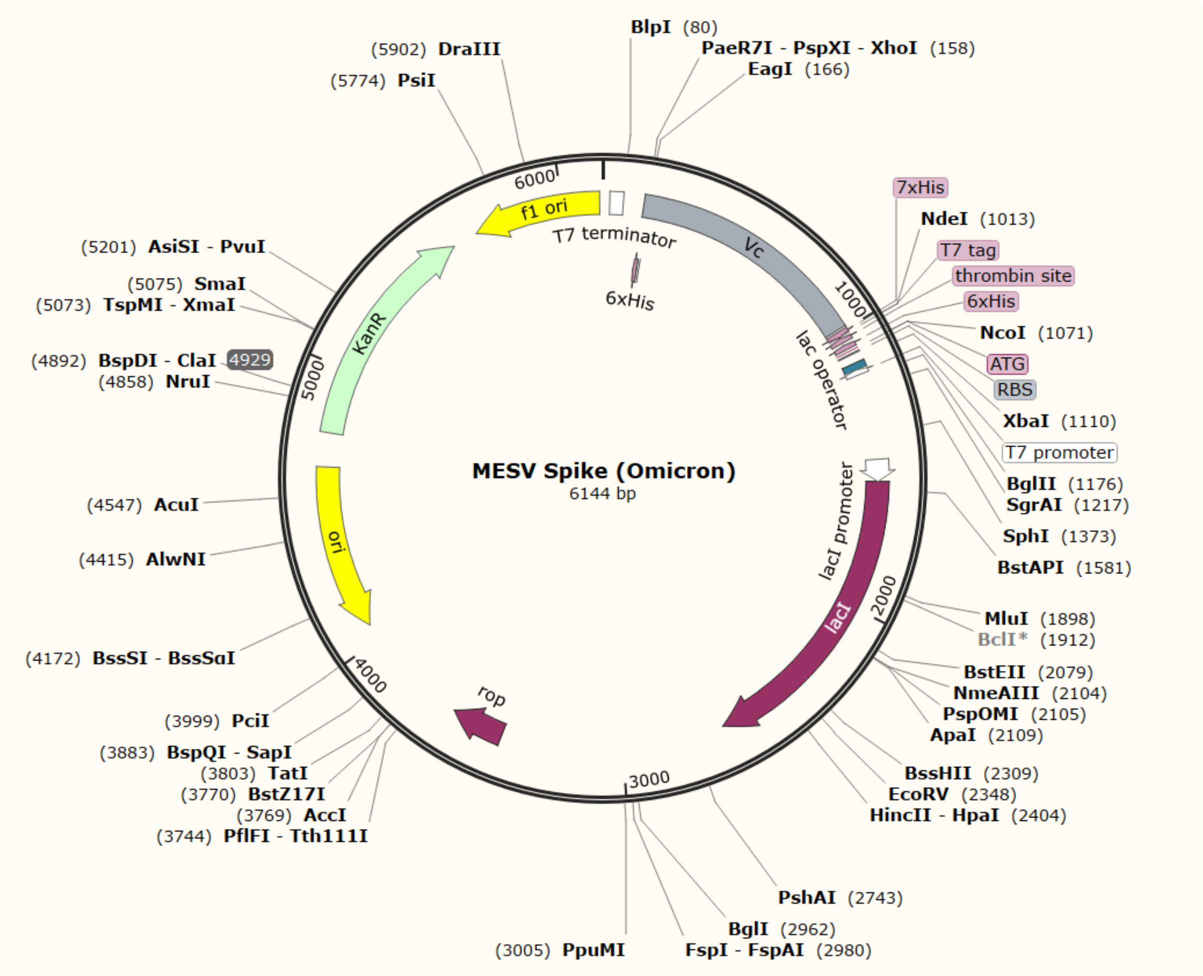


**Figure S2:** ***In silico*** **cloning of the final vaccine construct into pET28a (+) expression vector where the grey part indicates the coding gene for the vaccine surrounded between *Nde*I and *Eag*I in the vector backbone with OriC (yellow), promoter (magenta), and antibiotic resistance gene (green).**
